# Natural antibodies to polysaccharide capsules enable Kupffer cells to capture invading bacteria in the liver sinusoids

**DOI:** 10.1101/2024.04.26.591254

**Authors:** Xianbin Tian, Yanni Liu, Kun Zhu, Haoran An, Jie Feng, Linqi Zhang, Jing-Ren Zhang

**Affiliations:** Center for Infectious Biology, School of Medicine, Tsinghua University, Beijing, China; Tsinghua-Peking Center for Life Sciences, Tsinghua University, Beijing, China; Institute of Medical Technology, Peking University Health Science Center, Beijing, China; Department of Microbiology and Infectious Disease Center, Peking University Health Science Center, Beijing, China; State Key Laboratory of Microbial Resources, Institute of Microbiology, Chinese Academy of Sciences, Beijing, China

**Keywords:** Encapsulated bacteria, capsule, capsular polysaccharide, *Streptococcus pneumoniae*, *Klebsiella pneumoniae*, liver macrophage, Kupffer cell, complement and natural antibody

## Abstract

The interception of blood-borne bacteria in the liver defines the outcomes of invasive bacterial infections, but the mechanisms of this anti-bacterial immunity are largely speculative. This study shows that natural antibodies (nAbs) to capsules enable liver macrophage Kupffer cells (KCs) to rapidly capture and kill blood-borne encapsulated bacteria in mice. Affinity pulldown with serotype-10A capsular polysaccharides (CPS10A) of *S. pneumoniae* (*Spn*10A) led to the identification of CPS10A-binding nAbs in serum. The CPS10A-antibody interaction enabled KCs to capture *Spn*10A bacteria from the bloodstream, in part through complement receptors on KCs. The nAbs were found to recognize the β1-6-linked galactose branch of CPS10A, and similar moieties of serotype-39 *S. pneumoniae* and serotype-K50 *Klebsiella pneumoniae* capsules. More importantly, the nAbs empowered KCs to capture serotype-39 *S. pneumoniae* and serotype-K50 *K. pneumoniae* in the liver. Collectively, our data have revealed a highly effective immune function of nAb against encapsulated bacteria, and provided a proof of concept for treating septic bacterial diseases with monoclonal antibodies.

**Short summary:** Rapid capture of potentially harmful bacteria in blood by liver macrophages are vital for the blood sterility and health. This work reports how naturally occurring antibodies in the plasma enable macrophages to capture and kill blood-borne bacteria in the liver.

## INTRODUCTION

Invasive infections of encapsulated bacteria in sterile organ/tissue sites are a leading cause of human morbidity and mortality (Naghavi, 2022). Capsules, the outermost cellular structures of encapsulated bacteria, are known as “slippery” or antiphagocytic coat of encapsulated bacteria due to the resistance to phagocytic killing of host defense (Comstock and Kasper, 2006). Certain physical properties of the capsules (e.g., hyperviscosity and negative charge) hinder the recognition and binding by phagocytes to encapsulated bacteria (Brown and Gresham, 2012; Nahm and Katz, 2012; Taylor and Roberts, 2002). Virtually all bacterial capsules are composed of capsular polysaccharides (CPSs) (An et al., 2024; Whitfield et al., 2020). There are great inter- and intra-species variation in chemical structure and antigenicity of CPSs (An et al., 2024; Whitfield et al., 2020). Many bacteria produce a large number of capsule types (An et al., 2024). As examples, there are 103 capsular serotypes in *S. pneumoniae,* a leading cause of community-acquired pneumonia, blood infection and meningitis (Ganaie et al., 2023; Silva-Costa et al., 2024). Capsule types are associated with the virulence level of encapsulated bacteria as exemplified by the dominance of the low-numbered pneumococcal serotypes/serogroups in severe pneumonia in pre-antibiotic era (White, 1938) and of *Escherichia coli* K1 in neonatal meningitis (Robbins et al., 1974).

The liver is regarded as the vascular “firewall” against invading bacteria in the blood circulation (Balmer et al., 2014; Jenne and Kubes, 2013). The earlier studies demonstrated the importance of the liver in filtering blood bacteria is demonstrated by the dominant role of the organ in trapping blood-borne bacteria (Benacerraf et al., 1959; Brown et al., 1981; Gregory et al., 1996; Mackaness, 1962; Martin et al., 1949; Rogers, 1956). More recent investigations have shown the liver macrophages - Kupffer cells (KCs) as the major immune cells for the capture of commensal and potentially pathogenic bacteria (An et al., 2022; Broadley et al., 2016; Gola et al., 2021; Helmy et al., 2006; Huang et al., 2022; Kolaczkowska et al., 2015; Lee et al., 2010; Surewaard et al., 2016; Wong et al., 2013; Zeng et al., 2016). This functional dominance of KCs is accompanied by their exceptional representation in cell number among tissue resident macrophages, since KCs constitute approximately 90% of total tissue macrophages in humans and mice (Bilzer et al., 2006).

While the KC-mediated anti-bacterial immunity is vital for host blood sterility and health, it is largely unknown how KCs capture circulating bacteria with such the astonishing speed and capacity. The recent studies have highlighted the strict requirement of pathogen-recognition receptors for KC capture of blood-borne bacteria, likely due to the high shear force in the high-speed hepatic bloodstream (Kubes and Jenne, 2018). However, in contrast to the massive number of microbes that potentially enter the blood circulation, the full receptor arsenal for KC capture of invading bacteria is still a blackhole since only a few pathogen receptors on KCs have been identified to date. The complement receptor CRIg is the first known receptor for KC capture of complement C3-opsonized bacteria (Broadley et al., 2016; Helmy et al., 2006; Zeng et al., 2016). Our recent studies have shown that capsule types define bacterial survival in the bloodstream and thereby virulence by structure-dependent variability of capsules in escaping phagocytic capture of KCs in the liver (An et al., 2022; Huang et al., 2022). Based on their capacities of the immune evasion, encapsulated bacteria can be divided into the high-virulence (HV) and low-virulence (LV) categories. While the HV serotypes completely circumvent hepatic recognition, the LV counterparts are partially captured by KCs to various extents. The asialoglycoprotein receptor (ASGR) was identified as the first known receptor on KCs for pneumococcal LV serotype-7F and -14 capsules (An et al., 2022). However, it remains unknown how KCs recognize many other LV capsule types.

Natural antibodies (nAbs) are spontaneously produced by B-1 cells and marginal zone B cells without deliberate immunization (Galili et al., 1984; Martin et al., 2001; Springer et al., 1961). The majority of nAbs recognize polysaccharide antigens. Human and mouse plasma contain nAbs against phosphorylcholine of pneumococcal cell wall teichoic acid, which confers a modest level of protection against pneumococcal infection (Briles et al., 1981b; Haas et al., 2005). nAbs against the O127 lipopolysaccharide (LPS) of enteropathogenic *E. coli* promote pathogen capture by KCs (Zeng et al., 2018). However, the specific functions of many nAbs are largely speculative.

This study sought to identify host receptor(s) for the capsule of *S. pneumoniae* serotype-10A (*Spn*10A). *Spn*10A is a prevalent serotype in childhood invasive pneumococcal diseases (Silva-Costa et al., 2024), but displays a LV phenotype in mice (An et al., 2022). Natural antibodies were identified to specifically recognize serotype-10A capsular polysaccharide (CPS10A). We further revealed that this antibody-capsule binding interaction enables KCs to capture and eliminate blood-borne *Spn*10A in the liver. Broad function of the anti-CPS10A natural antibodies in host defense against other encapsulated bacteria was also validated.

## RESULTS

### Circulating *Spn*10A are captured by Kupffer cells

This study aimed to understand how KCs capture serotype-10A *S. pneumoniae* (*Spn*10A), one of the prevalent serotypes in the childhood pneumococcal diseases (Silva-Costa et al., 2024). We first verified the virulence phenotype of *Spn*10A isolates in mice. Contrast to 100% mortality of mice post intravenous (i.v.) infection with 10^6^ colony forming units (CFU) of the HV strains D39 (serotype 2) and TIGR4 (serotype 4), all mice survived infection of five selected *Spn*10A strains (**Fig. 1A**). Accordingly, mice infected with *Spn*10A bacteria became undetectable in the blood at 12 h post infection and remained undetectable ever since, whereas all of the D39- and TIGR4-infected mice showed severe bacteremia before falling to the infection (**Fig. 1B**). These data confirmed that pneumococcal serotype 10A is indeed a low-virulence (LV) serotype in mice.

**Figure 1.**
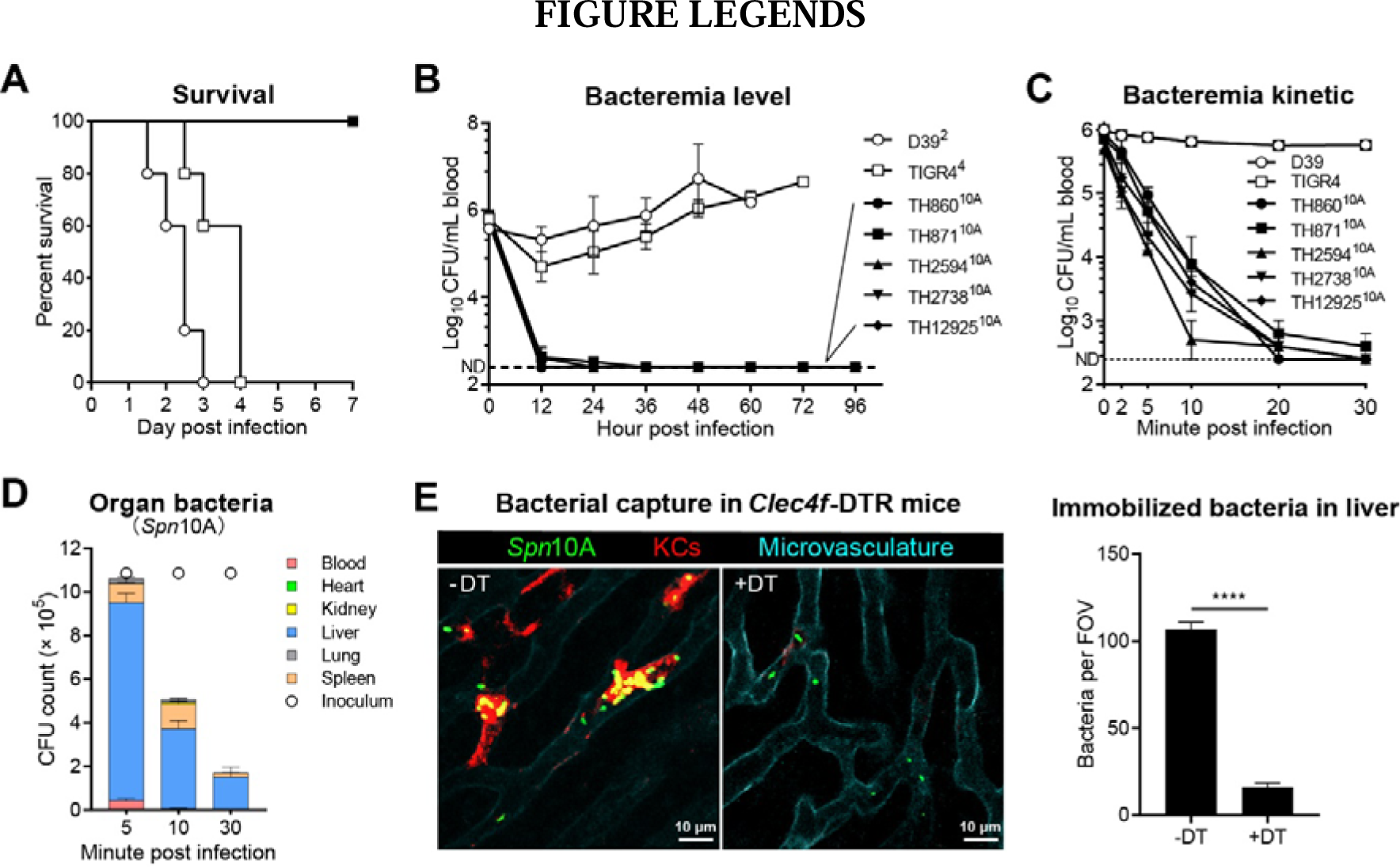
Capture of circulating *Spn*10A by liver KCs. **(A** and **B)** Survival rate (A) and bacteremia levels (B) of ICR mice intraperitoneally (i.p.) infected with 10^6^ CFU of D39, TIGR4 and five serotype-10A pneumococcal strains. Serotype of each strain is denoted with superscripted characters. n = 5. Dashed line represents detection limit of bacteremia. **(C)** Bacteremia kinetics of ICR mice in the first 30 min post intravenous (i.v.) infection with 10^6^ CFU of D39, TIGR4 and five serotype-10A pneumococcal strains. n = 3. **(D)** Bacterial distribution in the blood and organs of ICR mice at 5, 10 and 30 min post i.v. infection with 10^6^ CFU of TH860^10A^. n = 3. **(E)** Intravital microscopy (IVM) detection of pneumococcal capture in the liver sinusoids of *Clec4f*-DTR mice treated with (+DT) or without (-DT) diphtheria toxin post i.v. infection with 5 × 10^7^ CFU of TH860^10A^. Representative images (left) showed *S. pneumoniae* (green), KCs (red) and microvasculature (cyan) at 15 min post infection. n = 2. Quantitation of bacteria immobilized in liver in 5-10 random fields of view (FOVs) photographed at 10-15 min post infection is shown in the right. The processes of bacterial capture are demonstrated in Video 1. Unpaired *t* test (E) was performed. ****, P < 0.0001.

We further assessed the clearance process of *Spn*10A strains in the bloodstream. Opposite to the stable presence of D39 and TIGR4 in the bloodstream in the first 30 min of i.v. infection, all the five tested *Spn*10A strains rapidly disappeared from the circulation with extremely short 50% clearance time (CT_50_) of less than 3 min (**Fig. 1C**). We further characterized bacterial organ distribution at various time points post infection using representative strain TH860^10A^. The pneumococci were mostly trapped in the liver at 5 min (86% of detectable bacteria), 10 min (73% of detectable bacteria) and 30 min (69% of detectable bacteria) post inoculation (**Fig. 1D**). The spleen carried 8%, 22% and 28% of detectable bacteria at 5 min, 10 min and 30 min, respectively. There were marginal levels of bacteria in the lung, heart and kidney at all the time points. With the rapid disappearance of the bacteria from the circulation in the course of blood infection, these bacteria were effectively killed in the liver. The combined CFU values from the blood and five organs were similar to the inoculum at 5 min, but dramatically reduced to 47% and 7% of the inoculum at 10 min and 30 min, respectively. These results showed that *Spn*10A bacteria are rapidly captured and killed in the liver.

Based on the dominant role of liver macrophage Kupffer cells (KCs) in capturing the LV pneumococci (An et al., 2022), we visualized KC capture of *Spn*10A in the liver sinusoids by intravital microscopy (IVM) (**Fig. 1E**; **Video 1**). IVM imaging showed rapid tethering of circulating bacteria onto KCs post i.v. inoculation, but dramatical reduction of immobilized bacteria on KCs was observed with *Clec4f*-DTR mice treated with diphtheria toxin (DT). The KC-immobilized bacteria in each field of view (FOV) in KC-deficient mice was only 15% of what was observed at the vascular wall of the liver sinusoids in WT mice. These results demonstrated that KC is the dominant immune cell responsible for capturing *Spn*10A in the liver.

### Kupffer cells capture *Spn*10A by recognizing capsular polysaccharide

To understand how KCs capture *Spn*10A, we hypothesized that the macrophage recognizes the serotype-10 capsular polysaccharide (CPS10A), on the basis of our previous work with serotype-14 pneumococci (An et al., 2022). This possibility was tested by i.v. administration of purified CPS10A before i.v. bacterial inoculation. While CPS10A-treated mice showed stable bacteremia at various time points in the first 30 min post TH860^10A^ infection, similar treatment with purified capsular polysaccharide of serotype-14 (CPS14) did not show obvious impact on early clearance of CPS10A (**Fig. 2A**). Serotype-specific blocking of TH860^10A^ clearance with homologous but not heterologous CPS was also verified by CPS10A-dependent inhibition against *Spn*10A trapping in the liver. In contrast to dramatic disappearance of the *Spn*10A inoculum from the body of CPS14-treated mice at 30 min post i.v. inoculation with TH860^10A^, CPS10A-treated mice retained 97% of the inoculum at 30 min, the majority of which was located in the blood (**Fig. 2B**). These data indicated that KCs capture *Spn*10A by recognizing the capsular polysaccharide.

**Figure 2.**
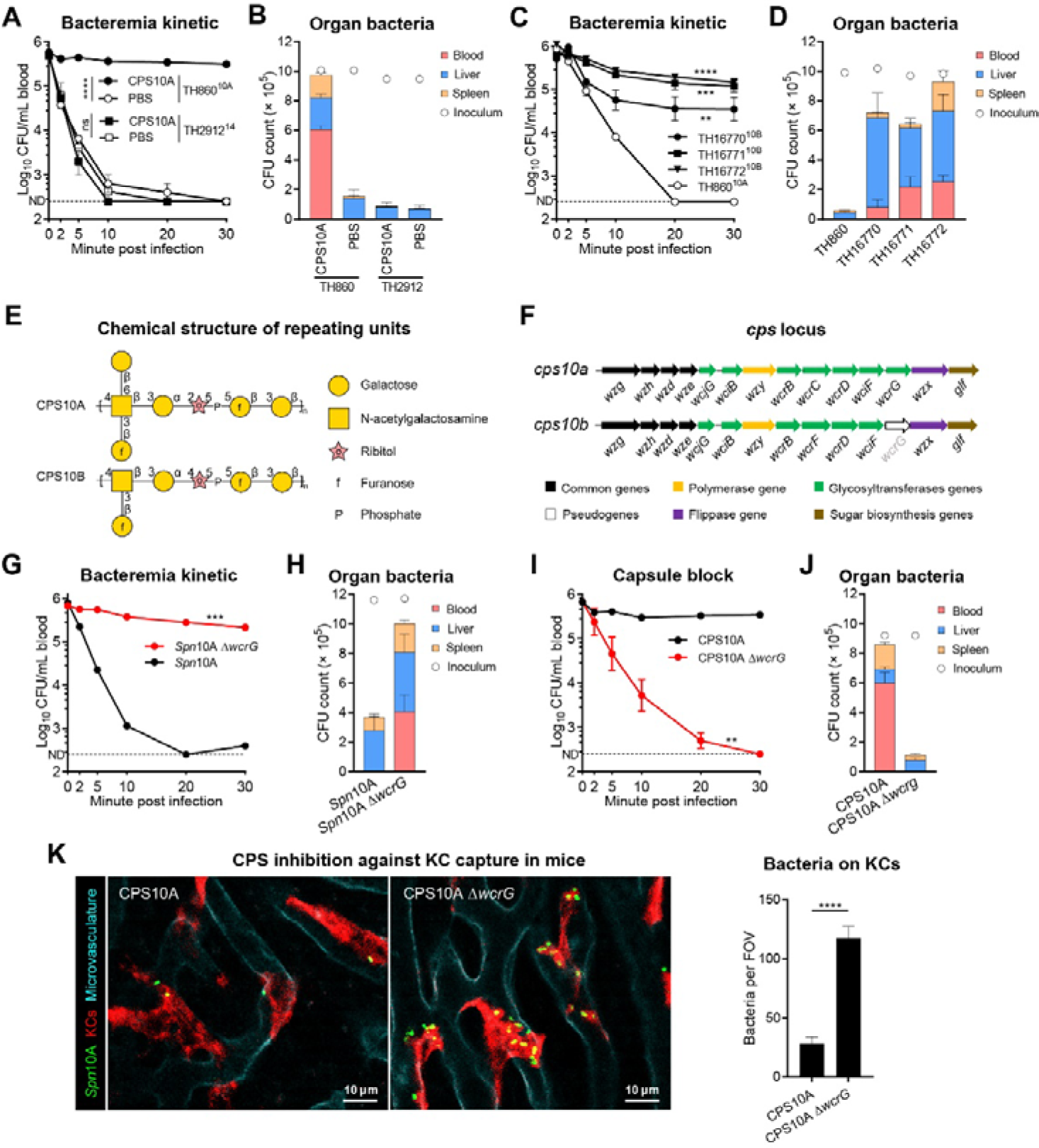
Capsular branches-dependent KC capture of circulating *Spn*10A. **(A** and **B)** Bacteremia kinetics in the first 30 min (A) and bacterial distribution at 30 min (B) post i.v. infection with 10^6^ CFU of TH860^10A^ or TH2912^14^ to ICR mice with i.v. treatment of 400 μg purified CPS10A or PBS 2 min before infection. n = 3. **(C** and **D)** Bacteremia kinetics in the first 30 min (C) and bacterial distribution at 30 min (D) of ICR mice post i.v. infection with 10^6^ CFU of TH860^10A^ or serotype-10B pneumococcal strains. n = 3. **(E)** Biochemical structure of repeating units of CPS10A and CPS10B. **(F)** Schematic representation of the *cps* loci of *Spn*10A and *Spn*10B. **(G** and **H)** Bacteremia kinetics in the first 30 min (G) and bacterial distribution at 30 min (H) of ICR mice post i.v. infection with 10^6^ CFU of TH860^10A^ or TH860^10A^ Δ*wcrG*. n = 3. **(I** and **J)** Bacteremia kinetics in the first 30 min (I) and bacterial distribution at 30 min (J) post i.v. infection with 10^6^ CFU of TH860^10A^ to ICR mice with i.v. treatment of 400 μg purified CPS10A or CPS10A Δ*wcrG* 2 min before infection. n = 3. **(K)** Representative IVM images of liver sinusoids (left) and quantitation of bacteria immobilized on KCs (right) of mice treated with 400 μg CPS10A or CPS10A Δ*wcrG* post i.v. infection with 5 × 10^7^ CFU of TH860^10A^. n = 2. The processes of bacterial capture were displayed in Video 2. Ordinary two-way ANOVA with Tukey’s multiple comparisons test (A, C, G and I) and unpaired *t* test (K) were performed. **, P < 0.01; ***, P < 0.001; ****, P < 0.0001; ns, not significant.

Since *Spn*10A belongs to serogroup 10 of *S. pneumoniae*, which contains five serotypes with similar capsular structures (Geno et al., 2015), we further assessed early clearance of three serotype-10B strains in our collection (TH16770, TH16771 and TH16772). Surprisingly, all the three strains showed much slower clearance than serotype-10A strains (**Fig. 2C**). Consistently, the CFUs of the serotype-10B strains recovered from the blood, liver and spleen at 30 min were significantly higher than that of TH860^10A^ (**Fig. 2D**). This result suggested that serotype-10B CPS does not bind to the host receptor(s) that recognizes CPS10A.

Serotypes 10A and 10B are highly similar in chemical structure (**Fig. 2E**) and setting of biosynthesis genes (**Fig. 2F**). However, CPS10A structurally differs from CPS10B in two aspects: a β1-6-linked galactopyranose (Gal*p*) branch (absent in CPS10B) and a α1-2 linkage between galactose and ribitol-5-phosphate (α1-4 linkage in CPS10B). We thus tested if the Gal*p* branch in CPS10A is responsible for the serotype-specific KC capture of *Spn*10A by mutating the *wcrG* gene in the *cps10a* locus of TH860, encoding the enzyme that adds β1-6-linked Gal*p* branches to GalNAc in the CPS10A repeating unit (Yang et al., 2011). *wcrG* in the *cps10b* locus is a pseudogene. Removing the branch moiety made TH860 much more resistant to the hepatic clearance. The CT_50_ of TH860 Δ*wcrG* mutant was elongated to 17.3 min from 1.1 min in the parental strain (**Fig. 2G**). The total bacteria detected from the blood, liver and spleen at 30 min post infection were significantly higher in mice infected with the mutant (**Fig. 2H**). Consistently, pre-treatment of mice with purified capsular polysaccharide from TH860 Δ*wcrG* (CPS10A Δ*wcrG*) did not hinder the shuffling of TH860 from the blood (**Fig. 2I**) to the liver (**Fig. 2J**) as compared with the intact CPS10A. IVM imaging showed five-fold more KC-immobilized bacteria in the mutant CPS-treated mice as compared with mice pre-treated with the normal capsular polysaccharide (**Fig. 2K**; **Video 2**). These data indicated that the β1-6-linked Gal*p* branch on CPS10A is recognized by an uncharacterized receptor(s) for KC capture of serotype-10A pneumococci.

### Serum antibodies specifically bind to the capsule of *Spn*10A

KCs may capture pathogens by membrane receptors directly or by soluble pattern recognition molecules indirectly (Kubes and Jenne, 2018). To determine whether KC-mediated capture of *Spn*10A relies on soluble component(s) in serum, we detected binding interactions between isolated KCs and bacteria in *in vitro* system. The visualization of KC binding showed that only by addition of mouse serum can KCs capture *Spn*10A and the inhibition was achieved by CPS10A but not CPS10A Δ*wcrG* (**Fig. 3A**). These data indicate that KCs capture *Spn*10A through recognizing β1-6-linked Gal*p* branch on CPS10A in the presence of serum.

**Figure 3.**
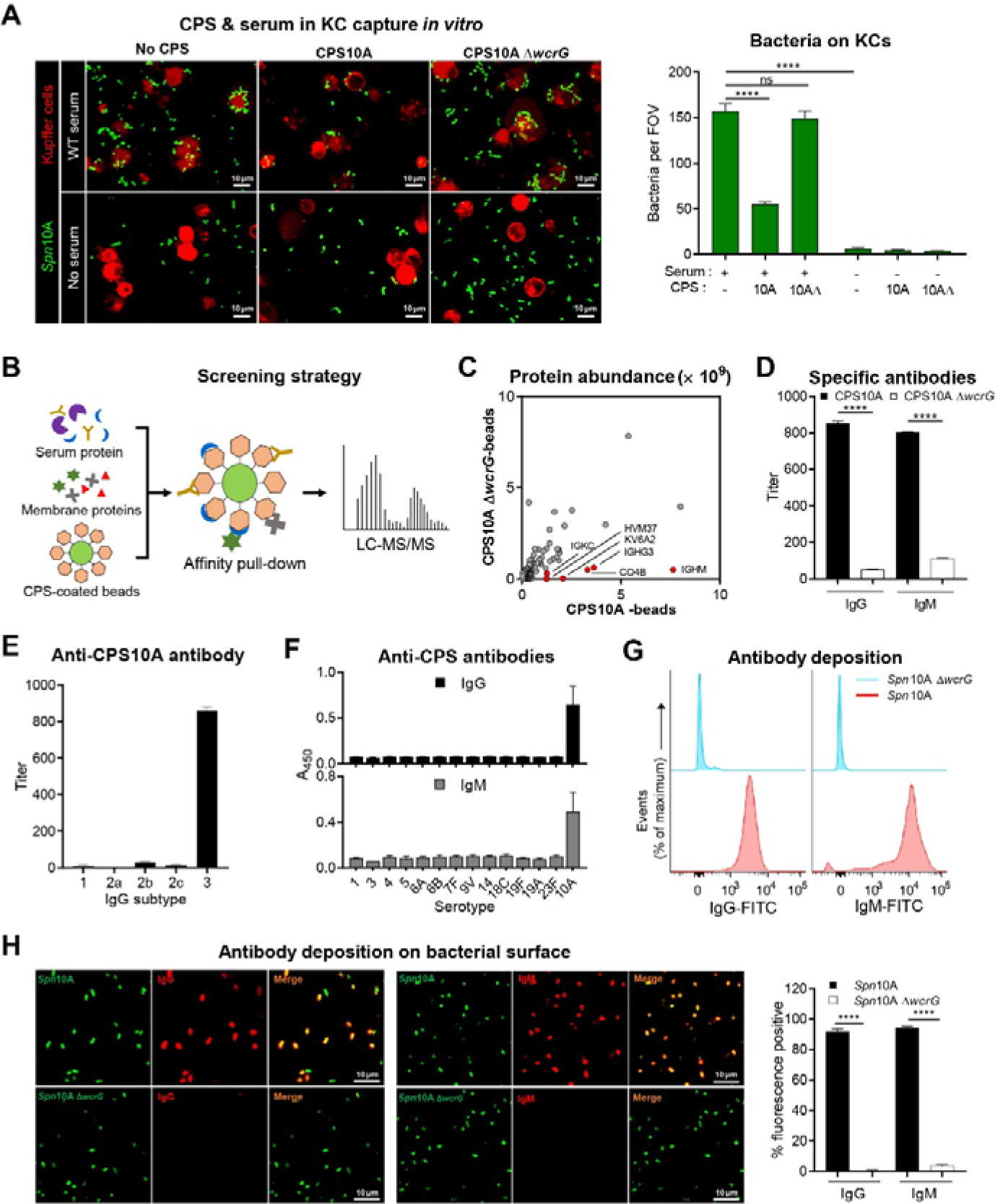
Serum antibodies specifically bind to the capsule of *Spn*10A. **(A)** Representative confocal images (left) and quantitative analysis (right) showing serotype-10A pneumococcal binding to isolated mouse KCs. The KCs were incubated with TH860 supplemented with or without 10% serum and 100 μg/ml CPS10A or CPS10A Δ*wcrG*. n = 6. **(B)** Schematic illustration of strategy for screening CPS10A-binding proteins. **(C)** Plot of proteins significantly enriched by CPS10A-coated beads. The top six enriched proteins compared to that of CPS10A Δ*wcrG*-coated beads are labelled in red. n = 3. The detailed results are listed in Table S1. **(D)** ELISA detection of IgG and IgM to CPS10A or CPS10A Δ*wcrG* in WT mouse serum. n = 3. **(E)** ELISA detection of IgG subtypes of anti-CPS10A IgG in mouse serum. n = 3. **(F)** ELISA detection of IgG and IgM to different pneumococcal CPSs in mouse serum. n = 3. **(G)** Flow cytometry detection of the IgG and IgM deposition on TH860^10A^ or TH860^10A^ Δ*wcrG* surface after incubation with mouse serum. **(H)** Representative confocal images showing the deposition of IgG (left) or IgM (middle) on TH860^10A^ or TH860^10A^ Δ*wcrG*. Quantitative of the percentage of TH860^10A^ or TH860^10A^ Δ*wcrG* with deposition of IgG or IgM was shown in the right. Picture with > 100 bacterial cells were chosen for quantitation. n = 5. Ordinary two-way ANOVA with Sidak’s multiple comparisons test (A, D and H) was performed. ****, P < 0.0001.

To identify host factor(s) that promotes KC capture of *Spn*10A, we performed an affinity pull-down of CPS10A-binding protein(s) in mouse serum and membrane proteins extracted from mouse nonparenchymal cells using CPS10A- and CPS10A Δ*wcrG*-coated beads (**Fig. 3B**). Proteomics analysis of the resulting proteins revealed that 37 proteins were at least two-fold enriched by CPS10A-coated beads (**Fig. 3C**; **Table S1**). With the consideration of protein abundance and enrichment fold, immunoglobulin heavy constant mu (IGHM), ig gamma-3 chain C region (IGHG3), complement C4-B (CO4B), ig kappa chain V-VI region XRPC 24 (KV6A2), ig heavy chain V region X44 (HVM37) and immunoglobulin kappa constant (IGKC) were among the most enriched proteins (**Fig. 3C**). Collectively, these hits represented various fragments of antibodies. In agreement, ELISA test revealed more CPS10A-reactive IgG and IgM in mouse serum than antibodies bound to CPS10A Δ*wcrG* (**Fig. 3D**). In addition, the CPS10A-reactive IgG antibodies almost exclusively belonged to the IgG3 subtype (**Fig. 3E**), which is coincidence with our mass spectrometry data. We further determined the capsule-reactive antibodies in mouse serum with purified CPSs from 13 additional serotypes of *S. pneumoniae* that are covered by the 13-valent pneumococcal polysaccharide conjugate vaccine (PCV13) (Briles et al., 2019). The binding interaction was not detected in any of these serotypes beyond CPS10A (**Fig. 3F**). This result excluded the possibility that the CPS10A-binding antibodies might be resulted from the well-known binding of pneumococcal cell wall phosphocholine to natural antibodies (Briles et al., 1981b). We also detected abundant anti-CPS10A antibodies in both male and female mice with different genetic backgrounds (**Figs. S1A** and **S1B**). Through flow cytometry, we detected IgG and IgM in serum deposited on *Spn*10A while no positive signals on surface of *Spn*10A Δ*wcrG* were observed (**Fig. 3G**). This was further confirmed by microscopy that more than 95% of *Spn*10A but no *Spn*10A Δ*wcrG* were covered by IgG and IgM in serum (**Fig. 3H**). Taken together, the data demonstrate that serotype-10A capsule is recognized by anti-CPS10A antibodies in serum.

### Anti-capsule antibodies enable KCs to capture *Spn*10A

To determine whether anti-CPS10A antibodies contribute to hepatic trapping of serotype-10A *S. pneumoniae*, we assessed early clearance of TH860^10A^ in μMT mice, which lack mature B cells and antibodies. The antibody-deficient mice normally cleared *Spn*14 as wide-type (WT) control in the first 30 min post infection, but completely failed to shuffle TH860^10A^ from the bloodstream (**Fig. 4A**) to the liver (**Fig. 4B**). Similar results were obtained with other four *Spn*10A strains (**Figs. S2A** and **S2B**). To verify the specific immunity of antibodies against *Spn*10A, we tested if mouse serum and purified immunoglobulins could rescue the immune deficiency of μMT mice in clearing *Spn*10A. Pre-instillation of mouse serum in μMT mice 5 min prior to i.v. bacterial inoculation led to a dose-dependent enhancement in early clearance of *Spn*10A (**Fig. 4C**). In a similar manner, the clearance of TH860^10A^ in μMT mice was restored by pre-administration of purified serum IgG (**Figs. 4D**) or IgM (**Fig. 4E**). These results showed that serotype-specific plasma antibodies are essential for hepatic clearance of blood-borne serotype-10A *S. pneumoniae*.

**Figure 4.**
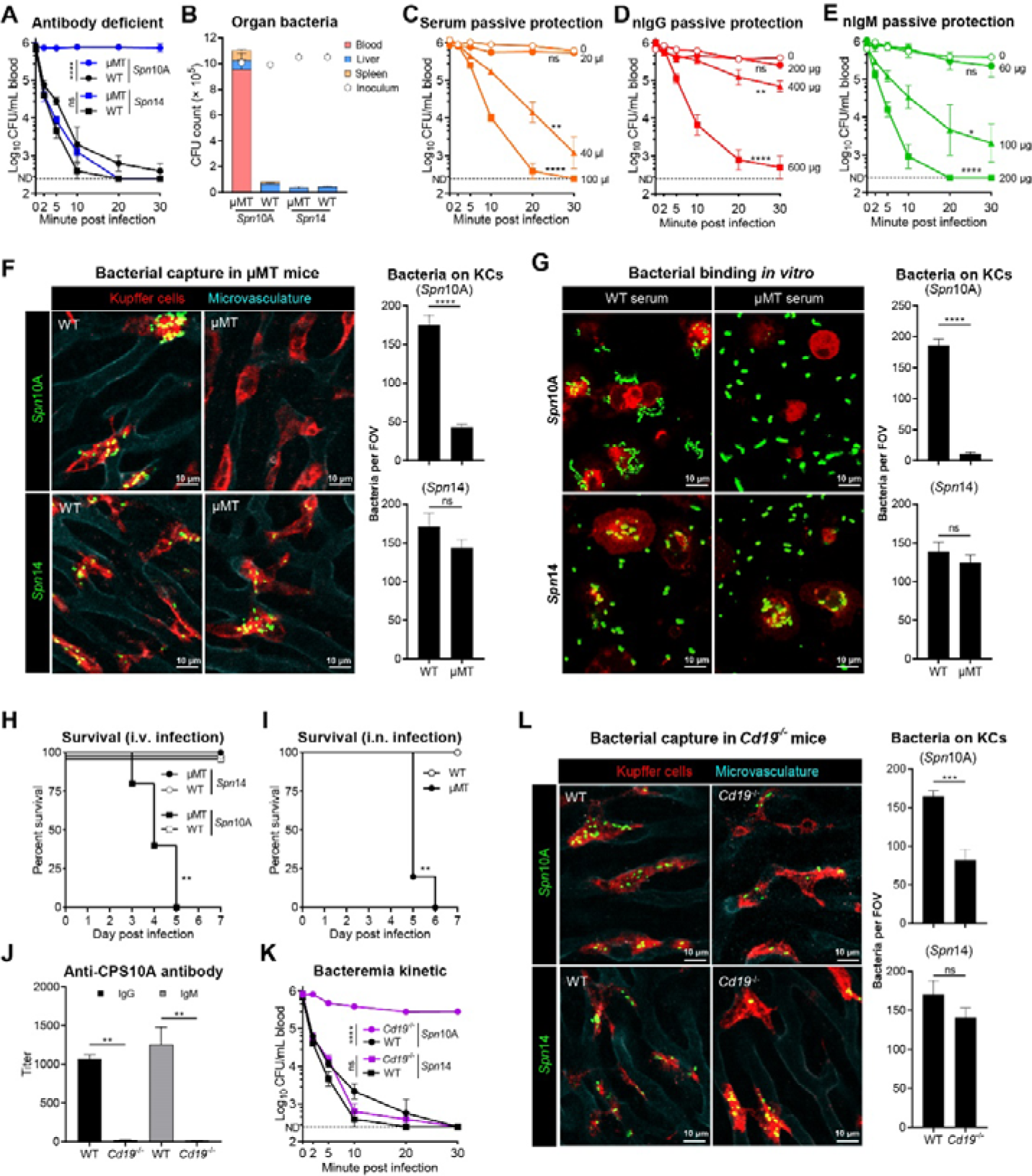
Clearance of *Spn*10A mediated by anti-CPS10A natural antibodies. **(A** and **B)** Bacteremia kinetics in the first 30 min (A) and bacterial distribution of WT or μMT mice at 30 min (B) post i.v. infection with 10^6^ CFU of TH860^10A^ or TH2912^14^. n = 3. **(C**-**E)** Bacteremia kinetics in the first 30 min post i.v. infection with 10^6^ CFU of TH860^10A^ to μMT mice with i.v. injection of 0, 20, 40 or 100 μl WT mouse serum (C), 0, 200, 400 or 600 μg purified natural IgG (D) or 0, 60, 100 or 200 μg purified natural IgM (E) 2 min before infection. n = 3. **(F)** Representative IVM images of liver sinusoids (left) and quantitation of bacteria immobilized on KCs (right) of WT or μMT mice post i.v. infection with 5 × 10^7^ CFU of TH860^10A^ or TH2912^14^. The processes of bacterial capture were displayed in Video 3 and Video 4. n = 2. **(G)** Representative confocal images showing TH860^10A^ or TH2912^14^ binding to isolated mouse KCs supplemented with 20% serum of WT or μMT mice. Quantitative of the KC-tethered bacteria was shown in the right. n = 5. **(H)** Survival rate of WT or μMT mice that were i.v. infected with 10^7^ CFUs of TH860^10A^ or TH2912^14^. n = 5. **(I)** Survival rate of WT or μMT mice that were i.n. infected with 10^7^ CFUs of TH860^10A^. n = 5. **(J)** ELISA detection of IgG and IgM to CPS10A in serum of WT and *Cd19*^-/-^ mice. n = 3. **(K)** Bacteremia kinetics of WT or *Cd19*^-/-^ mice in the first 30 min post i.v. infection 10^6^ CFU of TH860^10A^ or TH2912^14^. n = 3. **(L)** Representative IVM images of liver sinusoids (left) and quantitation of bacteria immobilized on KCs (right) of WT or *Cd19*^-/-^ mice post i.v. infection with 5 × 10^7^ CFU of TH860^10A^ or TH2912^14^. n = 2. The processes of bacterial capture are demonstrated in Video 4 and Video 5. Ordinary two-way ANOVA with Tukey’s (A, C, D, E and K) or Sidak’s (J) multiple comparisons test, unpaired *t* test (F, G and L) and log-rank test (H and I) were performed. *, P < 0.05; **, P < 0.01; ***, P < 0.001; ****, P < 0.0001; ns, not significant.

The immune function of anti-CPS10A antibodies was further manifested by antibody-dependent hepatic capture of TH860^10A^ via IVM imaging of the liver sinusoids. In contrast to effective capture of TH860^10A^ by KCs of WT mice, μMT mice displayed significant reduction in KC-associated pneumococci (**Fig. 4F**; **Video 3**). However, *Spn*14 bacteria were similarly captured by KCs between μMT and WT mice. We also verified the antibody-driven host-pathogen interaction under the *in vitro* conditions. Primary mouse KCs abundantly bound to TH860^10A^ bacteria in the presence of WT mouse serum, but KCs exhibited a marginal level of bacterial binding when being co-incubated with of μMT mouse serum (**Fig. 4G**). In sharp contrast, *Spn*14 bacteria were abundantly tethered to KCs regardless serum source, which agrees with the ASGR-mediated capture of *Spn*14 by KCs (An et al., 2022). Finally, septic infection experiments showed that the antibody-mediated bacterial capture in the liver is essential for host survival against serotype-10A *S. pneumoniae.* While all the tested WT mice survived infection with TH860^10A^ pneumococci that were inoculated via either i.v. route (**Fig. 4H**) or intranasal route (**Fig. 4I**), both the infection regiments resulted in 100% mortality of μMT mice. However, the antibody-deficient mice fully survived against i.v. infection with *Spn*14 pneumococci (**Fig. 4H**). These experiments demonstrated the essential role of antibody-mediated capsule recognition in serotype-specific host defense against serotype-10A *S. pneumoniae*.

The presence of relatively high level of anti-CPS10A antibodies in the serum of naive mice strongly suggested the nature of natural antibody (nAb) (Kearney et al., 2015). This possibility was tested using *Cd19*^-/-^ mice that lack B-1a cells, the major source of natural antibodies (Baumgarth et al., 2005; Rickert et al., 1995). The anti-CPS10A antibodies were barely detected in serum of *Cd19*^-/-^ mice (**Fig. 4J**). Accordingly, *Cd19*^-/-^ mice were found to display severe impairment in the clearance of *Spn*10A but not *Spn*14 bacteria (**Fig. 4K**). Specific impairment in hepatic capture of *Spn*10A but not *Spn*14 bacteria was also observed in *Cd19*^-/-^ mice by IVM imaging (**Fig. 4L**; **Video 4** and **Video 5**). These data allowed us to conclude that the anti-CPS10A antibodies belong to natural antibodies.

### The complement system is required for nAb-mediated hepatic clearance of *Spn*10A

To explore the mechanism of nAb-mediated clearance of *Spn*10A, we assessed potential involvement of antibody receptors using mice lacking the known IgM receptors (FCAMR *-Fcamr*^-/-^ and FCMR - *Fcmr*^-/-^) or four IgG receptors FcγRI-IV (FcRα null). However, none of these mice showed obvious deficiency in early clearance of *Spn*10A as compared to WT mice (**Fig. S3A**), indicating that KCs do not use these known Fc receptors for binding to nAb-opsonized *Spn*10A. In the light of our earlier observation that complement components C1q, C1s, C1r, C4b and C3 were abundantly enriched by CPS10A-coated beads (**Figure. S3B**; **Table S1**), we hypothesized that KCs rely on the complement system to capture nAb-opsonized *Spn*10A. Because the classical pathway of the complement system is known to mediate antibody-driven phagocytosis (Zipfel and Skerka, 2009), we tested potential contribution of complement proteins C1q and C3, two key components of the classical pathway. Clearance of *Spn*10A were significantly delayed in both *C1qa*^-/-^ and *C3*^-/-^ mice, although the former displayed more severe phenotype (CT_50,_ 12 min) than *C3*^-/-^ mice (CT_50,_ 1.6 min), as compared to that in WT mice (CT_50,_ 0.9 min) (**Fig. 5A**). In the same fashion, *C1qa*^-/-^ and *C3*^-/-^ mice showed lower levels of liver-trapped bacteria (**Fig. 5B**). The defect of *C1qa*^-/-^ and *C3*^-/-^ mice in bacterial clearance was not due to potential difference in nAb production because these mice showed a similar level of anti-CPS10A nAbs as WT mice (**Fig. S4**). These experiments indicated that anti-CPS10A nAbs enable KCs to capture *Spn*10A, at least in part by activating complement-mediated phagocytosis.

**Figure 5.**
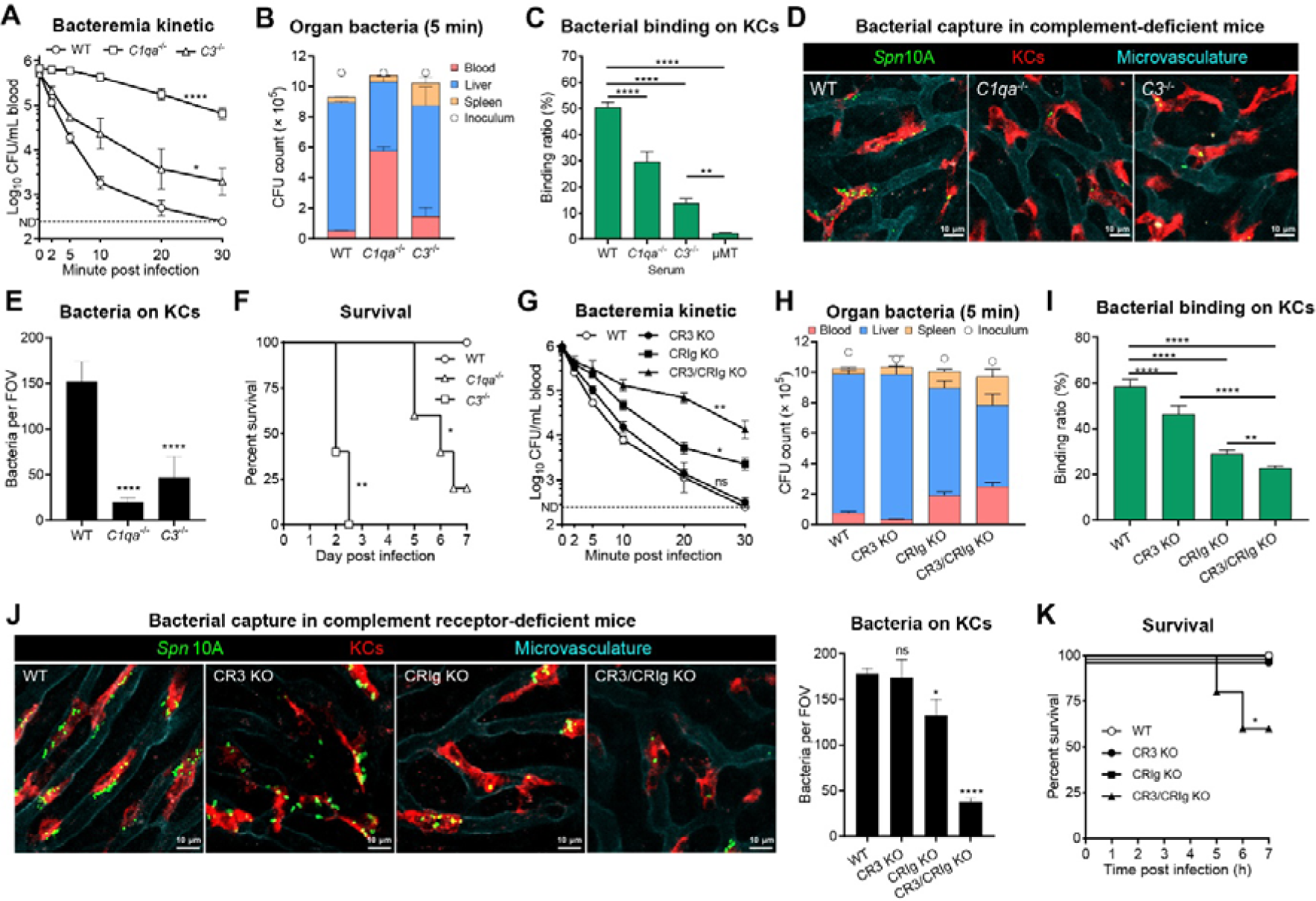
Indispensable role of complement system in natural antibodies-mediated *Spn*10A capture. **(A** and **B)** Bacteremia kinetics in the first 30 min (A) and bacterial distribution at 5 min (B) of WT, *C1qa*^-/-^ or *C3*^-/-^ mice post i.v. infection with 10^6^ CFU of TH860^10A^. n = 3. **(C)** TH860^10A^ binding to isolated KCs from WT, *C1qa*^-/-^, *C3*^-/-^ and μMT mice with incubation of 10% WT serum. n = 3. **(D** and **E)** Representative IVM images of liver sinusoids (D) and quantitation of bacteria immobilized on KCs (E) of WT, *C1qa*^-/-^ or *C3*^-/-^ mice post i.v. infection with 5 × 10^7^ CFU of TH860^10A^. n = 2. The processes of bacterial capture are demonstrated in Video 6. **(F)** Survival rate of WT, *C1qa*^-/-^ or *C3*^-/-^ mice that were infected with 10^8^ CFU of TH860^10A^. n = 10. **(G** and **H)** Bacteremia kinetics in the first 30 min (G) and bacterial distribution at 5 min (H) of WT, CR3 KO (*Itgam*^-/-^), CRIg KO (*Vsig4*^-/-^) or CR3/CRIg KO (*Itgam*^-/-^ *Vsig4*^-/-^) mice post i.v. infection with 10^6^ CFU of TH860^10A^. n = 3. **(I)** TH860^10A^ binding to isolated KCs from WT, CR3 KO, CRIg KO or CR3/CRIg KO mice with incubation of 10% WT serum. n = 3. **(J)** Representative IVM images of liver sinusoids (left) and quantitation of bacteria immobilized on KCs (right) of WT, CR3 KO, CRIg KO or CR3/CRIg KO mice post i.v. infection with 5 × 10^7^ CFU of TH860^10A^. n = 2. The processes of bacterial capture are demonstrated in Video 7. **(K)** Survival rate of WT, CR3 KO, CRIg KO or CR3/CRIg KO mice that were infected with 10^8^ CFUs of TH860^10A^. n = 5. Ordinary two-way ANOVA with Tukey’s multiple comparisons test (A and G), one-way ANOVA with Tukey’s multiple comparisons test (C, E, I and J) and log-rank test (F and K) were performed. *, P < 0.05; **, P < 0.01; ***, P < 0.001; ****, P < 0.0001; ns, not significant.

We further validated the contribution of the complement system to KC capture of *Spn*10A under the *in vitro* conditions using sera from *C1qa*^-/-^ and *C3*^-/-^ mice. Both sera from *C1qa*^-/-^ mice and *C3*^-/-^ mice were less able to promote KC binding to *Spn*10A (**Fig. 5C**). However, these sera displayed relatively milder phenotypes than that from μMT mice, suggesting that nAbs promote bacterial clearance in the liver by an uncharacterized complement-independent mechanism(s). In a similar manner, IVM imaging showed that KCs of *C1qa*^-/-^ and *C3*^-/-^ mice are significantly defective in capturing *Spn*10A in the liver sinusoids (**Figs. 5D** and **5E**; **Video 6**). Consistently, *C1qa*^-/-^ and *C3*^-/-^ mice succumbed to otherwise nonlethal infection of *Spn*10A (**Fig. 5F**). These results fully demonstrated that the complement classical pathway is vital for nAb-mediated capture of *Spn*10A by KCs.

To determine how KCs capture nAb/C3-opsonized pneumococci, we tested the role of CR3 and CRIg, two complement receptors that are significantly expressed on KCs (Helmy et al., 2006). While CR3-deficient (*Itgam*^-/-^) mice did not show any obvious defect in early clearance of *Spn*10A, CRIg-deficient (*Vsig4*^-/-^) mice displayed a modest phenotype (**Fig. 5G**). However, simultaneous loss of both the receptors (CR3/CRIg KO) yielded a more significant impact on bacterial clearance (CT_50_, 4.7 min). In a similar manner, mice with the combined absence of CR3 and CRIg (CR3/CRIg KO) showed the most severed impairment in hepatic capture of *Spn*10A than the mice lacking either the receptor (**Fig. 5H**). The functional redundancy of CR3 and CRIg in mediating KC capture of nAb/C3-opsonized pneumococci was also verified with primary KCs. KCs from CR3/CRIg KO mice displayed lower capacity of bacterial binding than those from the single receptor-deficient mice (**Fig. 5I**). Likewise, IVM imaging also revealed more severe deficiency with CR3/CRIg KO mice than CR3- or CRIg-deficient mice in KC capture of *Spn*10A in the liver sinusoids (**Fig. 5J**; **Video 7**). Accordingly, CR3/CRIg KO mice were more susceptible to infection of *Spn*10A (**Fig. 5K**). Collectively, our data demonstrated that the complement classical pathway and C3 receptors are indispensable in KC capture of nAb/C3-opsonized *Spn*10A.

### Anti-CPS10A nAbs recognize other Gal*p* branch-containing capsules

Our mass spectrometry data (**Fig. 3B**; **Table S1**) revealed significant enrichment of variable regions of multiple monoclonal antibodies that have been previously reported, including XRPC 24 (X24) and XRPC 44 (X44) that are known to recognize β1-6 galactan (Rudikoff et al., 1973). Since CPS10A contains β1-6 galactose, we tested whether these monoclonal antibodies react with CPS10A by generating recombinant IgM and IgG3 forms of X24 and X44 based on the additional sequence information available in the UniProt databases. ELISA results showed specific binding to CPS10A by the IgM and IgG3 forms of X24 and X44 (**Fig. 6A**). IgM form of both antibodies had relatively higher levels of binding affinity to CPS10A than the IgG3 counterpart, which might be due to different valences of IgM (pentamer) and IgG3 (monomer).

**Figure 6.**
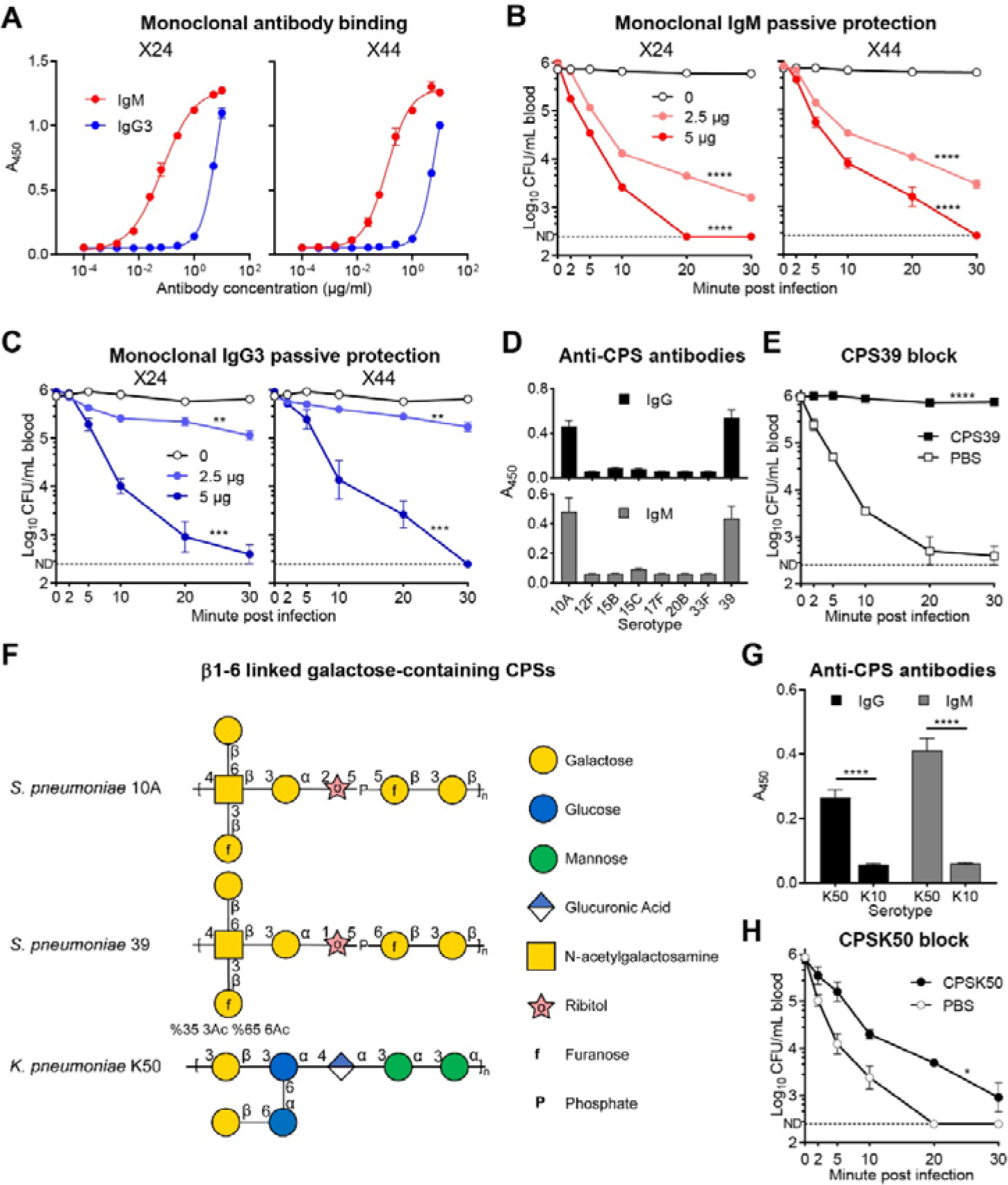
Recognition of other Gal*p* branch-containing capsules by anti-CPS10A natural antibodies. **(A)** ELISA detection of X24-IgM, -IgG3 (left) and X44-IgM, -IgG3 (right) binding to CPS10A. n = 3. **(B** and **C)** Bacteremia kinetics in the first 30 min post i.v. infection with 10^6^ CFU of TH860^10A^ to μMT mice with i.v. treatment of 0, 2.5 and 5 μg X24 (left) or X44 (right) -IgM (B) and -IgG3 (C) 2 min before infection. n = 3. **(D)** ELISA detection of antibodies to CPSs with galactose branch in murine serum. n = 6. **(E)** Bacteremia kinetics in the first 30 min post i.v. infection with 10^6^ CFU of TH860^10A^ to mice with i.v. treatment of 400 μg purified CPS39 or PBS 2 min before infection. n = 3. **(F)** Biochemical structure of repeating units of CPS10A, CPS39 and *Klebsiella pneumoniae* K50 CPS. **(G)** ELISA detection of IgG and IgM to *K. pneumoniae* K50 and K10 CPS in serum of WT mice. n = 6. **(H)** Bacteremia kinetics in the first 30 min post i.v. infection with 10^6^ CFU of TH860^10A^ to mice with i.v. treatment of 800 μg purified *K. pneumoniae* K50 CPS (CPSK50) or PBS 2 min before infection. n = 3. Ordinary two-way ANOVA with Tukey’s multiple comparisons test (B, C, E and H) and one-way ANOVA with Tukey’s multiple comparisons test (G) were performed. *, P < 0.05; **, P < 0.01; ***, P < 0.001; ****, P < 0.0001.

Importantly, both the IgM form (**Fig. 6B**) and IgG3 form (**Fig. 6C**) of the antibodies significantly accelerated early clearance of serotype-10A pneumococci in antibody-deficient μMT mice in a dose-dependent manner when they were i.v. inoculated 2 min prior to bacterial infection. However, the IgM antibodies were in general more effective than the IgG3 counterparts when both the antibody forms were used at the same concentrations. While the IgM antibodies were still effective at the dose of 2.5 µg (**Fig. 6B**), but the IgG3 proteins showed only marginal impact on pneumococcal clearance at this dose (**Fig. 6C**). In the context of functional dependence of nAbs on the complement system (**Fig. 5A**), this functional difference could be caused by higher potency of IgM antibodies in C3 activation (Hajishengallis et al., 2017). These results indicated that anti-CPS10A nAbs and previously described anti-β1-6 galactan antibodies belong to the same pool of anti-polysaccharide nAbs in mice.

Many pneumococcal serotypes contain galactose branches in the capsular repeating units (Geno et al., 2015), including α1-2 Gal*p* in serotypes 15B, 15C and 33F, α1-4 Gal*p* in serotype 17F, β1-3 Gal*p* in serotype 12F, β1-4 Gal*f* in serotype 20B and β1-6 Gal*p* in serotype 39 (**Fig. S5**). We thus determined if anti-CPS10A nAbs react with any of these galactose branch-containing capsules. ELISA test revealed significant binding between purified serum IgG and IgM and serotype-39 capsular polysaccharide (CPS39), but not obvious antibody binding was detected with the other six serotypes (**Fig. 6D**). To determine whether the same natural antibodies bind to both CPS10A and CPS39, we performed *in vivo* competitive blocking of *Spn*10A clearance with purified CPS39. Pre-treatment with CPS39 remarkably inhibited early clearance of *Spn*10A (**Fig. 6E**), which was at a similar extent to what was observed with CPS10A (**Fig. 2A**). This finding is consistent with the presence of a β1-6-linked Gal*p* branch and an identical glycan composition in both CPS10A and CPS39, although there are significant differences between the two serotypes in glycan linkage and acetylation (Petersen et al., 2014) (**Fig. 6F**). Through additional search, we also realized that the capsule of serotype-K50 *Klebsiella pneumoniae* (CPSK50) also contain the terminal β1-6-linked Gal*p* (Altman and Dutton, 1983) (**Fig. 6F**). ELISA revealed significant binding of CPSK50 to purified serum IgG and IgM (**Fig. 6G**). Consistently, pre-treatment of mice with CPSK50 significantly blocked early clearance of *Spn*10A (**Fig. 6H**). These results uncovered broad recognition of bacterial capsules by circulating nAbs.

### Anti-capsule nAbs enhance hepatic clearance of multiple encapsulated bacteria

Since anti-CPS10A nAbs also recognized CPS39, we determined whether the nAbs played a role in early clearance of serotype-39 pneumococci (*Spn*39). In contrast to severe impairment of antibody-deficient μMT mice in clearing *Spn*10A (**Fig. 4A**), *Spn*39 bacteria were similarly cleared between the mutant and WT mice (**Fig. 7A**). This result suggested that *Spn*39 bacteria are cleared by nAb-mediated and additional redundant mechanisms or the nAb-mediated *in vitro* CPS39 binding does not operate *in vivo*. A previous study has shown that nAbs significantly enhance hepatic clearance of enteropathogenic *Escherichia coli* cells in the absence of the complement system (*C3*^-/-^ mice) (Zeng et al., 2018). We thus tested early clearance of *Spn*39 in *C3*^-/-^ μMT mice lacking both antibodies and C3. The double knockout mice displayed severe deficiency in shuffling *Spn*39 bacteria from the blood (**Fig. 7A**) to the liver (**Fig. 7B**), indicating the anti-capsule nAbs and complement system represent two unique immune mechanisms against *Spn*39. In line with this conclusion, *C3*^-/-^ mice exhibited a significantly but partial defect in early clearance of this bacterium. The impact of this dual immune mechanisms on hepatic clearance of *Spn*39 was also observed by IVM imaging of the liver sinusoids. *Spn*39 bacteria were abundantly captured by KCs of *C3*^-/-^ and μMT mice, but completely bypassed the hepatic trapping in *C3*^-/-^ μMT mice (**Figs. 7C** and **7D**; **Video 8**).

**Figure 7.**
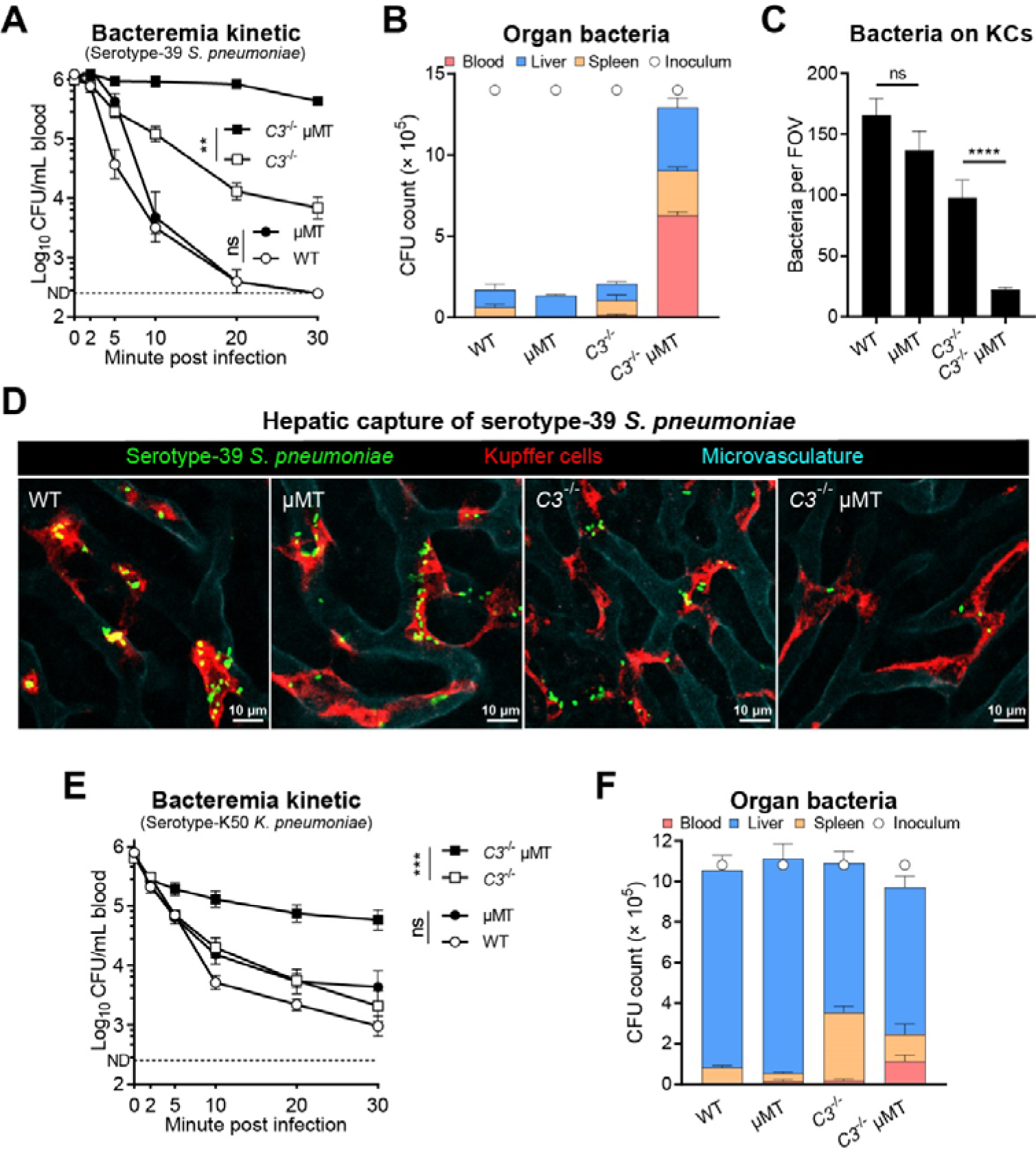
Clearance of Gal*p* branch-containing encapsulated bacteria mediated by natural antibodies. **(A** and **B)** Bacteremia kinetics in the first 30 min (A) and bacterial distribution of WT, μMT, *C3*^-/-^ or *C3*^-/-^ μMT mice at 30 min (B) post i.v. infection with 10^6^ CFU of TH16827^39^. n = 3. **(C** and **D)** Quantitation of bacteria immobilized on KCs (C) and representative IVM images of liver sinusoids (D) of WT, μMT, *C3*^-/-^ or *C3*^-/-^ μMT mice post i.v. infection with 5 × 10^7^ CFU of TH16827^39^. The processes of bacterial capture are demonstrated in Video 8. n = 2. **(E** and **F)** Bacteremia kinetics in the first 30 min (E) and bacterial distribution of WT, μMT, *C3*^-/-^ or *C3*^-/-^ μMT mice at 30 min (F) post i.v. infection with 10^6^ CFU of TH17033^K50^. n = 6. Ordinary two-way ANOVA with Tukey’s multiple comparisons test (A and E) and one-way ANOVA with Tukey’s multiple comparisons test (C) were performed. **, P < 0.01; ***, P < 0.001; ****, P < 0.0001; ns, not significant.

We finally determined potential contribution of nAb-mediated capsule binding to host immunity against serotype-K50 *K*. *pneumoniae,* a major Gram-negative pathogen in hospital-acquired infections (Paczosa and Mecsas, 2016). As observed with *Spn*39 (**Fig. 7A**), the early clearance of *K*. *pneumoniae* was marginally affected by the loss of either antibody (in μMT mice) or C3 protein (in *C3*^-/-^ mice) (**Fig. 7E**). However, simultaneous absence of antibody production and C3 protein in *C3*^-/-^ μMT mice led to more pronounced defect in clearing blood-borne *Spn*39. Consistently, the liver of *C3*^-/-^ μMT mice showed a relatively lower level of bacterial burden as compare with those of *C3*^-/-^ or μMT mice (**Fig. 7F**). Therefore, anti-CPS10A nAbs contribute to clearance of bacteria with β1-6-linked Gal*p* branch in capsule. Taken together, these data had revealed that anti-capsule natural antibodies can confer broad protection against multiple encapsulated bacteria.

## DISCUSSION

While receptor-mediated bacterial capture by KCs in the liver is vital for host blood sterility and health, only a few such receptors are known. As a matter of fact, the scavenger receptor ASGR is the only capsule receptor that mediates KC capture of encapsulated bacteria, which are frequently associated with bloodstream infections and septic deaths (Naghavi, 2022). With the lead of our recent findings that KCs are able to recognize the LV capsule types (An et al., 2022; Huang et al., 2022), this study has shown that the anti-β1-6 galactan nAbs recognize the capsules of serotype-10A and -39 *S. pneumoniae* and serotype-K50 *K. pneumoniae*. More importantly, the nAb-capsule interactions enable KCs to capture these pathogens in the liver. Our data have highlighted that nAbs serves as potent capsule receptors for KC capture of potentially virulent encapsulated bacteria. To the best of our knowledge, this is the first report that explicitly define nAbs as capsule receptors that mediate hepatic clearance of invading encapsulated bacteria.

### Naturally antibodies are the serotype-specific receptors for multiple bacterial capsules

Natural antibodies are known to fulfil various immune functions, such as clearance of apoptotic cells, regulation of B cell immune responses and broad defense against invading pathogens (New et al., 2016). Natural antibodies are known to recognize broad spectrum of bacteria, fungi, viruses and parasites (Choi and Baumgarth, 2008; Sarden et al., 2022; Subramaniam et al., 2010; Yilmaz et al., 2014; Zeng et al., 2018). The nAbs recognizing pneumococcal cell wall phosphocholine is the prototype of anti-microbial nAbs, mainly owing to the early availability of anti-phosphocholine hybridoma antibodies (Andres et al., 1981; Gearhart et al., 1981). More importantly, the anti-phosphocholine antibodies are protective against the bloodstream infections of *S. pneumoniae* (Arai et al., 2011; Briles et al., 1981a; Briles et al., 1992; Briles et al., 1981b). Zeng *et al*. have recently reported that nAbs against O127 lipopolysaccharide (LPS) of enteropathogenic *E. coli* drive pathogen capture by KCs (Zeng et al., 2018). To the best of our knowledge, no capsule-specific nAbs have been explicitly documented. This study revealed the abundant presence of naturally occurring antibodies toward the capsules of important human pathogens - *S. pneumoniae* and *K. pneumoniae*.

Our finding of IgM and IgG3 as natural antibodies to bacterial capsules is in line with the dominance of these two antibody subtypes in serum (Palma et al., 2018). In addition, most of nAbs in human and mouse recognize core glycans on glycoproteins and glycolipids (New et al., 2016). Finally, nAbs are mainly produced by B-1 cells and marginal zone B cells, which distinguish antigen-induced antibodies produced by B-2 cells (Martin et al., 2001). Accordingly, the anti-capsule nAbs were barely detectable in *Cd19*^-/-^ mice that lack B-1a cells.

### Natural antibodies to CPSs enable KCs to capture circulating bacteria

KCs are extremely capable of capturing the LV capsule types of encapsulated bacteria (An et al., 2022; Huang et al., 2022), but ASGR is the only characterized capsule receptor. In this study, we first verified the LV phenotype of serotype-10A *S. pneumoniae* isolates and their rapid capture by KCs in the liver. We further demonstrated CPS10A as a specific ligand for KC recognition by competitive blocking of bacterial clearance with purified CSP10A. These lines of information led to the discovery of nAbs as CSP10A-binding proteins by affinity pulldown and protein mass spectrometry. The immune function of anti-CPS10A nAbs was defined by significant impairment of antibody-deficient mice (μMT and *Cd19*^-/-^ mice) in early blood clearance and hepatic interception of serotype-10A *S. pneumoniae*. Intravital microscopy imaging allowed us to visualize and quantify the antibody-dependent immune action of KCs in capturing fast-flowing bacteria in the liver sinusoids. This nAb-mediated KC capture of serotype-10A *S. pneumoniae* is reminiscent of our recent finding that vaccine-induced antibodies enable KCs to capture the otherwise “uncatchable” HV capsule types of *S. pneumoniae* (Wang et al., 2023). Zeng *et al*. have recently shown that nAbs makes KCs able to capture enteropathogenic *E. coli* by recognizing O127 LPS (Zeng et al., 2018). While the authors did not explicitly characterize the contribution of potential nAb-capsule interactions to the phenotype, it appears that the capsule-mediate binding is, at least partially, responsible for the KC capture of Xen-14 *E. coli* since the strain utilizes the same polysaccharide repeating unit for LPS and capsule biosynthesis. To this end, it is reasonable to believe that nAbs are important for hepatic clearance of invading bacteria.

### Anti-β1-6 galactan nAbs broadly recognize β1-6-linked Gal*p* on multiple CPSs

nAbs are long known to confer protective immunity against microbial infections, but the precise mechanisms of nAb-mediated protection are mostly speculative at the molecular level. A major hurdle against comprehensively studying nAb-mediated anti-bacterial immunity is the lack of pathogen-specific antigens recognized by natural antibodies and appropriate model systems. Built on our recent finding that liver macrophages effectively capture serotype 10A and other LV capsule types of *S. pneumoniae* in a receptor-dependent manner in mouse blood infection model (An et al., 2022; Huang et al., 2022), we identified CPS10A-binding nAbs by the affinity pulldown approach and subsequently verified the functional importance of these antibodies in activating KC capture of serotype-10A pneumococci in the liver sinusoids. Extended experiments also revealed a rather broad immune recognition of the capsules from additional encapsulated Gram-positive (serotype-39 *S. pneumoniae*) and Gram-negative (serotype-K50 *K. pneumoniae*) bacteria. More importantly, nAb-mediated capsule recognition activates pathogen capture and killing by KCs in the liver. Our mutagenesis work in the capsule biosynthesis gene locus of serotype-10A *S. pneumoniae* initially demonstrates the functional importance of the β1-6-linked Gal*p* on CPS10A in KC capture of the bacteria. Our later affinity pulldown with CPS10A-coated beads dramatically enriched nAbs that resemble the various regions of the β1-6-linked galactan-binding nAbs, particularly XRPC-24 and XRPC-44. Consistent with the nature of nAbs, the purified preparations of these antibodies from mouse serum were identified to be the IgM and IgG3 forms. Interestingly, both the IgM and IgG3 forms of the nAbs rescued the functional defect of antibody-deficient mice in clearing serotype-10A *S. pneumoniae*.

The XRPC-24 and XRPC-44 monoclonal antibodies were originally identified as the IgA form from myeloma in BALB/c mice following i.p. injection of mineral oil, and were found to bind with β1-6-linked galactotetraose (Potter et al., 1972; Rudikoff et al., 1973). In the following two decades, the amino acid sequences of these antibodies and other β1-6-linked galactan-binding nAbs were determined (Rao et al., 1979; Rudikoff et al., 1984; Rudikoff et al., 1983; Rudikoff et al., 1980). This information served as importance clues for functional characterization of the CPS10A-binding nAbs in this work. We generated recombinant forms of XRPC-24 and XRPC-44 in both the IgM and IgG3 antibody types, and demonstrated that these antibodies behave in the same way as the CPS10A-binding nAbs isolated from mouse serum. This and other lines of information have provided convincing evidences that the CPS10A-binding nAbs represent the same antibody repertoire as XRPC-24 and XRPC-44, which recognize the β1-6-linked Gal*p* epitope.

The realization of antigenic nature of the CPS10A-binding nAbs made it possible to search for other encapsulated bacteria with the β1-6-linked Gal*p* moiety on their capsule polysaccharide repeats. This led to our subsequent investigation of serotype-39 *S. pneumoniae* and serotype-K50 *K. pneumoniae*, which were found to contain the β1-6-linked Gal*p* moiety in the repeat units of the capsules in the Carbohydrate Structure Database (CSDB) (Toukach and Egorova, 2016). To another level, subsequent experiments confirmed the functional contribution of the CPS10A-binding nAbs to hepatic clearance of serotype-39 *S. pneumoniae* and serotype-K50 *K. pneumoniae*. While it remains to be determined if the CPS10A-binding nAbs recognize additional polysaccharides, the β1-6-linked Gal*p* moiety is a common component of *E. coli* LPS O antigens (e.g., O46, O64, O124, O131, O134, O164 and O171) (Liu et al., 2020). In the context of the previous finding that nAbs to the O127 antigen of enteropathogenic *E. coli* activate KC capture of the circulating bacteria (Zeng et al., 2018), it is reasonable to believe that the CPS10A-binding nAbs represent a broad type of pattern-recognition receptors that enable KCs to capture invading bacteria and perhaps other microbes bearing the β1-6-linked Gal*p* moiety.

### Natural antibodies mediate KCs capture of encapsulated bacteria by complement-dependent and -independent mechanisms

Antibodies typically mediate microbial elimination through binding interactions with Fc receptors on immune cells once their variable regions bind to specific antigens on the surface of microbes (West and Kemper, 2022). This principle is manifested by our recent finding that vaccine-induced IgG antibodies immobilize the otherwise liver-resistant capsule types of *S. pneumoniae* to the surface of liver sinusoidal endothelia cells, leading to effective bacterial capture and killing (Wang et al., 2023). However, our limited investigation has not identified specific Fc receptors that convey the nAb-opsonized bacteria to KCs. The IgG3-mediated KC capture of serotype-10A *S. pneumoniae* fully operated in mice lacking all of the four known phagocytic Fc receptors for IgG (FcγRI-IV). Likewise, the deletion of two know IgM receptors in mice did not compromise the IgM-mediated KC capture of the bacterium. While it remains to be determined if other uncharacterized Fc receptor(s) on KCs is responsible for physical engaging the nAb-opsonized pneumococci, the available data support an important role of the complement system in executing the nAb-initiated bacterial capture by KCs. This is manifested by delayed bacterial clearance in C3-deficient mice.

Antigen binding of IgG3 and IgM has been shown to activate complement C3 via the C1q-dependent classic pathway and thereby mediate internalization of immune complex through complement receptors (Azeredo da Silveira et al., 2002; Han et al., 2001; Ogden et al., 2005; Saylor et al., 2010; Weinstein et al., 2015). The current data support an important role of complement receptors CRIg and CR3 on KCs in capturing antibody/C3-opsonized *S. pneumoniae*. Based on the more pronounced phenotype in CRIg KO mice than CR3-deficient animals, it appears that CRIg plays a more prominent role in immobilizing nAb/C3-opsonized *S. pneumoniae*. C1q is known to activate C3 via the classical pathway by binding to the Fc region of antibodies once antibody-antigen complexes form (West and Kemper, 2022). The functional deficiency of C1qa KO mice in the nAb-mediated bacterial capture in the liver indicates the complement classical pathway is involved in the nAb-initiated C3 activation on CPS10A and other nAb-binding capsules. This conclusion is also supported by the CPS10A-mediated pulldown of various components of the classical pathway (e.g., C1q, C1s, C1r, C4b and C3). However, we noticed that C1q-deficient mice displayed a more severe impairment than C3-deficient the in the nAb-mediated *Spn*10A clearance. This information points to a complement-independent mechanism behind the nAb-mediated hepatic anti-bacterial immunity. This notion is supported by the phenotypic differences among mouse lines lacking C3 and antibodies. The clearance of *Spn*10A was partially retarded in C3-deficient mice, but completely lost in μMT mice.

In this context, we envision that natural IgM and IgG3 antibodies activate KCs via distinct pathways. On one hand, IgM nAb-mediated bacterial capture by KCs more relies on the complement system since pneumococcal polysaccharide vaccine-induced IgM antibodies fully depend on complement receptors CRIg and CR3 to mediate KC capture of high-virulence *S. pneumoniae* (Wang et al., 2023). It is difficult to image that natural IgM antibodies would take on a completely different path to initiate bacterial capture by KCs. On the other hand, IgG3 may functionally engage an uncharacterized Fc receptor(s) on KCs. The early study reported the existence of an mouse IgG3-specific receptor other than known FcγRs on J774A.1 cell line (Diamond and Yelton, 1981). Although mouse FcγRI was reported to bind with mouse IgG3-coated sheep erythrocytes with low affinity (Gavin et al., 1998), mouse IgG3-mediated phagocytosis of *Cryptococcus neoformans* was not affected in the absence of FcγRs (Saylor et al., 2010). Integrin β1 was regarded as part of a mouse IgG3 receptor, despite no evidence for a direct interaction (Hawk et al., 2019). The nAb-mediated KC capture of *Spn*10A may provide a new model for identifying novel IgG3 Fc receptor(s), since the nAbs alone confer a strong functional phenotype - shuffling *Spn*10A from the bloodstream to the liver sinusoids.

### The anti-capsule natural antibodies may have profound implications for the control of encapsulated bacterial diseases

Invasive infections by encapsulated bacteria represent a major threat to public health because many encapsulated bacteria are resistant to many or nearly all of the clinical antimicrobials, such as *Acinetobacter baumannii, E. coli* and *K. pneumoniae*. As manifested by the broad recognition of the β1-6-linked Gal*p* moiety on multiple capsules by the anti-CPS10A antibodies, broad-spectrum monoclonal antibodies against certain common antigenic epitopes of capsular polysaccharides may be developed as an alternative to antimicrobials for future treatment of drug-resistant encapsulated pathogens when the routine treatment practiced become obsolete.

## MATERIALS AND METHODS

### Bacterial strains, cultivation and genetic manipulation

Bacterial strains used in this research are listed in Table S2. Pneumococci were grown in Todd-Hewitt broth supplemented with 0.5% yeast extract (THY) or tryptic soy agar plates with 5% defibrated sheep blood at 37°C with 5% CO_2_ as described (Lu et al., 2006). For animal infection, bacteria were grown to OD_620 nm_ of 0.5 to 0.55, stored as frozen stocks, and diluted to desirable concentrations in Ringer’s solutions as inoculum. The precise concentration of each inoculum was determined by plating the inoculum immediately before infection.

Pneumococcal Δ*wcrG* mutant was generated in serotype-10A strain TH860 by natural transformation as described (Liu et al., 2019). TH860 was transformed with PCR fragments of the *rpsL1* allele to gain streptomycin resistance (Sung et al., 2001). The *wcrG* of TH860 was then replaced with Janus Cassette segments containing a *rpsL^+^* allele by using homologous recombination, followed by counterselection with the 1500-bp homologous flanking regions of *wcrG*. The relevant primers (Pr18745-Pr18750) are listed in Table S3.

### Mouse infection

All infection were performed in 6-8-week-old female C57/BL6 or ICR mice obtained from Vital River (Beijing, China). All experiments on mice were performed according to the animal protocols approved by the Institutional Animal Care and Use Committees at Tsinghua University. All genetic modified mice were maintained in C57/BL6 background. *C3*^-/-^ and μMT mice were obtained from the Jackson Laboratory. CR3 KO (*Itgam*^-/-^) mice were generated by CRISPR/Cas9 as described (Wang et al., 2023). CRIg KO *(Vsig4*^-/-^) mice were acquired from Genentech. *C1qa*^-/-^, *Cd19*^-/-^ and *Fcamr*^-/-^ mice were obtained from GemPharmatech (Nanjing, China). *Fcmr*^-/-^ mice were purchased from the Model Organisms Center (Shanghai, China). Mice that lack all mouse FcγR α-chain genes (FcRα null mice) were generally provided by professor Jeffrey V. Ravetch (Smith et al., 2012). *Clec4f*-DTR mice were generated and generally provided by professor Martin Guilliams (Scott et al., 2016). Depletion of KCs in *Clec4f*-DTR mice was achieved by i.v. injection of 25 ng/g diphtheria toxin to mice 24 h prior to infection. The other mice with multiple genetic modifications were obtained by cross breeding.

Animal survival was monitored for 7 days or at a humane endpoint (body weight loss > 20%). Bacteremia level was assessed by retroorbital bleeding and CFU plating. For bacterial early clearance, bacteremia level was assessed at various time points post intravenous (i.v.) infection. The clearance half time (CT_50_) values were calculated using the formula CT_50_ = ln(1 - 50/plateau)/(- *K*), in which plateau and *K* were generated through one-phase association nonlinear regression analysis of clearance ratio of each strain using GraphPad Prism as described previously (An et al., 2022). Bacteria in different organs were enumerated by plating homogenates of the liver, spleen, lung, kidney and heart. To determine the factors that affect bacterial clearance, CPSs, serum and purified antibodies in volume of 100 μl were i.v. injected to mice 2 min before i.v. infection if necessary. The volumes of intraperitoneal (i.p.), i.v. and intranasal infection were 200 μl, 100 μl and 30 μl, respectively.

### Intravital microscopy (IVM)

IVM imaging of mouse liver was performed as described (An et al., 2023). Briefly, 10^8^ CFU of pneumococci were stained in 100 μl PBS containing 10 μg FITC (Sigma) in the dark for 30 min. The hepatic microvasculature and KCs were stained by i.v. injection of 2.5 μg AF594 anti-CD31 and AF647 anti-F4/80 antibodies, respectively, for 30 min before i.v. inoculation with 5 × 10^7^ CFU of FITC-labeled *S. pneumoniae*.

Images were acquired with Leica TCS SP8 confocal microscope using 10×/0.45 NA and 20×/0.80 NA HC PL APO objectives. The microscope was equipped with Acousto Optics without filters. Fluorescence signals were detected by photomultiplier tubes and hybrid photo detectors (600 × 600 pixels for time-lapse series and 1,024 × 1,024 pixels for photographs). Three laser excitation wavelengths (488, 585 and 635 nm) were employed by white light laser (1.5 mw, Laser kit WLL2, 470-670 nm). Real-time imaging was monitored for 1 min after infection. Five to ten random fields of view at 10-15 min were selected to calculate the bacteria number per field of view.

### CPS purification

Purification of pneumococcal CPSs was as described with modification (Lee et al., 2020). Briefly, pneumococci were grown in THY medium and lysed by sodium deoxycholate solution. The pH of the lysate was adjusted to 5.0 to precipitate most soluble proteins. After centrifugation, the supernatant containing CPS was subjected to ultrafiltration against 25 mM sodium acetate to remove cell wall polysaccharide. The CPS was then precipitated by 3% hexadecyl trimethyl ammonium bromide (CTAB) and resuspended in 1 M NaCl solution. Ethanol was sequentially added to the solution to a final concentration of 33% to remove impurities and 80% to precipitate pneumococcal CPSs. CPS of *K. pneumoniae* was extracted as described (Huang et al., 2022).

### Screening of CPS10A-binding proteins

The identification of CPS-binding proteins by affinity pulldown of CPS-coated beads to serum proteins was performed as described with modifications (An et al., 2023). Briefly, CPSs were incubated with 4-(4,6-dimethoxy-1,3,5-triazin-2-yl)-4-methylmorpholinium chloride (DMTMM, Sigma-Aldrich). The mixture was then ultrafiltered to remove extra DMTMM and mixed with carboxyl latex beads (2 μm; Invitrogen). The beads were blocked in PBS with 1% bovine serum albumin (BSA). The membrane proteins of liver non-parenchymal cells (NPCs) were purified using the Mem-PER Plus Membrane Protein Extraction Kit (Thermo Fisher Scientific)

To isolate CPS-binding proteins, 1 × 10^8^ CPS-coated beads were mixed with 100 μl freshly isolated mouse serum and 100 μg purified membrane proteins of liver NPCs in total of 500 μl PBS at 37°C for 1 h. The beads were washed and then heated at 96°C for 5 min to dissociate CPS-binding proteins. The dissociated proteins were then separated on SDS-PAGE gels to excise protein bands for in-gel digestion. The protein mass spectrometry and analysis were performed as described (An et al., 2022). Protein abundance was compared between the CPS10A-and CPS10A Δ*wcrG*-coated beads, and proteins with more than two-fold preference to CPS10A beads were regarded as CPS10A-binding candidates.

### *In vitro* bacterial binding to Kupffer cells

Kupffer cells were isolated by a collagenase-DNase digestion procedure as described (Li et al., 2014). Briefly, mouse liver was perfused from the portal vein with digestion buffer containing HBSS with 0.5 mg/ml collagenase IV (Sigma-Aldrich), 20 μg/ml DNase I (Roche) and 0.5 mM CaCl_2_ and then digested in digestion buffer for 30 min at 37°C. The homogenate of liver was filtered through 70-μm strainers and washed. The residue red blood cells were lysed by RBC lysis buffer (BioLegend), and hepatocytes were removed by centrifugation at 50 g for 1 min. The liver NPCs in the supernatant were collected.

Bacterial binding to mouse KCs was performed as described (An et al., 2022). The liver NPCs were seeded into 48-well plate in RPMI 1640 and incubated for 30 min at 37°C to enrich KCs. The adherent cells (KCs) were then incubated 1 × 10^5^ CFU of bacteria (MOI = 1), 10% mouse serum and 100 μg/ml CPS (if necessary) in each well. The plate was then centrifugated at 500 g for 5 min and incubated for 30 min at 37°C with 5% CO_2_. The number of unbound bacteria in culture supernatant were determined by plating. The KCs were lysed by sterile water to determine the number of bound bacteria. Bacterial binding ratio was equal to the ratio of the bound bacteria number to the total bacteria number after the incubation.

To visualize *in vitro* bacterial binding to KCs, the NPCs were blocked by 1% anti-mouse CD16/CD32 for 10 min and then stained with AF647 anti-F4/80 (1:200, BioLegend) for 20 min. The 8 × 10^5^ cells/chamber stained cells were seeded into 4-Chamber 35 mm glass bottom dishes with 20 mm microwell (Cellvis) and cultured as mentioned in 48-well plates. After non-adherent cells were removed, 2 × 10^6^ CFU of bacteria (MOI = 10), 20% mouse serum and 100 μg/ml CPS (if necessary) were added to each chamber in a total volume of 500 μl. The dishes were then centrifugated at 500 g for 5 min and incubated for 30 min at 37°C with 5% CO_2_. The images were acquired using the Leica TCS SP8 confocal microscope randomly and pictures with more than 100 bacteria were chosen for quantification.

### Enzyme-linked immunosorbent assay (ELISA)

CPS-specific antibody was quantified by ELISA as described (Tian et al., 2024). Briefly, 96-well plates were coated with CPSs and blocked by 5% non-fat dry milk (BD Difco). The mouse serum was serially diluted and added into CPS-coated wells for 2-hour incubation. The detection of anti-CPS antibodies in mouse serum was performed by incubation with HRP-conjugated goat anti-mouse IgG (1:2000, EasyBio) or HRP-conjugated goat anti-mouse IgM (1:2000, EasyBio). To determine the subtypes of mouse anti-CPS10A IgG, HRP-conjugated IgG1-, IgG2b-, IgG2c- and IgG3-specific antibodies (1:2000, Proteintech) were used. The absorbance at wavelength of 450 nm was detected after incubation with TMB substrate (TIANGEN) and termination. The titer was calculated as highest dilution degree with positive/negative (P/N) ratio > 2.

### Fluorescence imaging and flow cytometry detection of antibody deposition

Briefly, 10^8^ CFU of pneumococci were stained in 100 μl PBS containing 10 μg FITC (Thermo) or AF647-NHS ester (Invitrogen) for 30 min at room temperature (RT). The bacteria were washed three times with PBS, followed by incubation with 50 μl serum for 30 min at RT. After three-time wash with PBS, the bacteria were incubated with FITC-conjugated goat anti-mouse IgG (1:2,000, EasyBio) or FITC-conjugated goat anti-mouse IgM (1:2,000, Elabscience) for 1 h. FITC signals on bacteria were detected by flow cytometry. Fluorescence images were acquired using the Leica TCS SP8 confocal microscope.

### Antibody production and purification

The total natural IgG was purified from murine serum using Protein G resin (GenScript, China) according to the manufacturer’s instructions. The total natural IgM was purified from the flow through serum from Protein G resin using Protein L resin (GenScript, China).

Recombinant monoclonal antibodies were generated as described (Wang et al., 2023). The coding sequences of light chains and heavy chains of X24 and X44 were synthesized based on amino acid sequences (Uniprot: KV6A1_MOUSE, HVM37_MOUSE, KV6A2_MOUSE, HVM39_MOUSE) and cloned into vectors containing constant region for light chain (pTH16770) or constant region for heavy chain of murine IgG3 (pTH16771) as described (Wang et al., 2023). The coding sequences of heavy chains were fused with PCR-amplified J-chain (GenBank: AB644392.1) and constant region of heavy chain of murine IgM (CDS of GenBank: BC096667.1) and cloned into vector. The related primers and plasmids are listed in Tables S3 and S4. The antibodies were produced by co-transfection of the vectors containing light chains and heavy chains to human HEK293 suspension culture cells (Expi293F) and purified with Protein G resin (IgG) and Protein L (IgM) resin, respectively.

### Statistical analysis

All experiments were performed with at least two biological replicates. Statistical analyses were performed using GraphPad Prism software (8.3.0). The error bars depict the standard error of mean (SEM). The significance was determined as described in figure legend. A *P* value of < 0.05 was considered significant.

## ACKNOWLEDGEMENTS

We thank J. V. Ravetch (rockefeller University, USa) and F. li (Shanghai Jiao Tong University, china) for providing Fcγr-humanized mice, and M. Guilliams (Ghent University) for providing *Clec4f*-DTR mice. We thank the Tsinghua research platforms for assistance in animal experimentation (Laboratory Animal Research Center), flow cytometry and IVM imaging (Center for Cell Biology) and protein mass spectrometry (Center for Proteomics). This work was supported by national natural Science Foundation of China grants 82330071, 31820103001, and 31530082 (J.-R.Z.) and 32200143 (Y.L.); national Key research and Development Program of China grant 2023YFc2306301 (J.-R.Z.); and Tsinghua-Peking Joint Center for Life Sciences Postdoctoral Foundation (Y.L.).

## AUTHOR CONTRIBUTIONS

Author contributions: Conceptualization, X. Tian and J.-R. Zhang; experimentation, X. Tian, Y. Liu, and H. An; methodology, K. Zhu, J. Feng, and L. Zhang; data analysis, X. Tian, Y. Liu, and J.-R. Zhang; composition, X. Tian and J.-R. Zhang; funding, J.-R. Zhang and Y. Liu.

---

**Figure S1.**
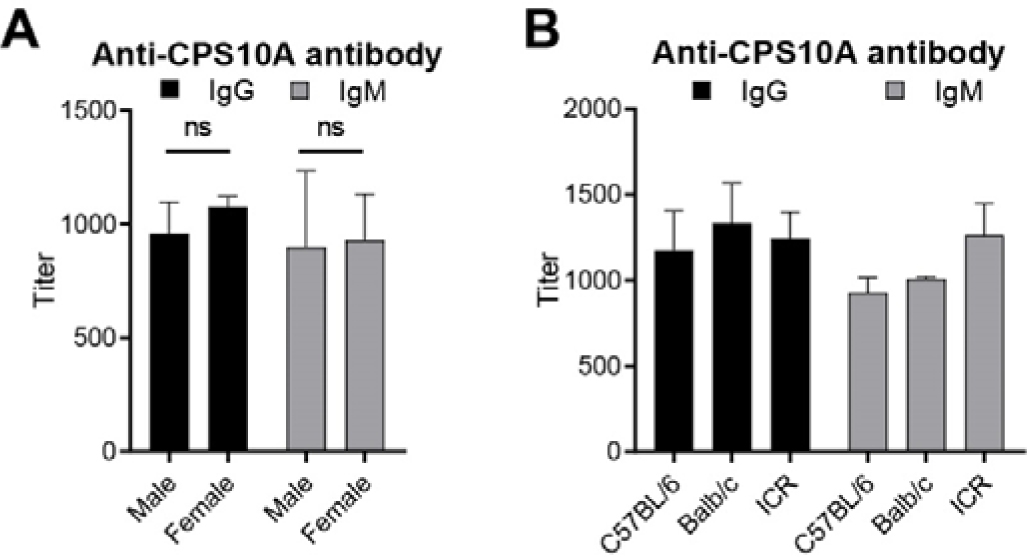
The wide presence of anti-CPS10A antibodies in mouse serum. **(A)** ELISA detection of IgG and IgM to CPS10A in serum of female and male mice. n = 5. **(B)** ELISA detection of serum IgG and IgM to CPS10A in C57BL/6, Balb/c or ICR mice. n = 3. Ordinary two-way ANOVA with Sidak’s multiple comparisons test (A) was performed. ns, not significant.

**Figure S2.**
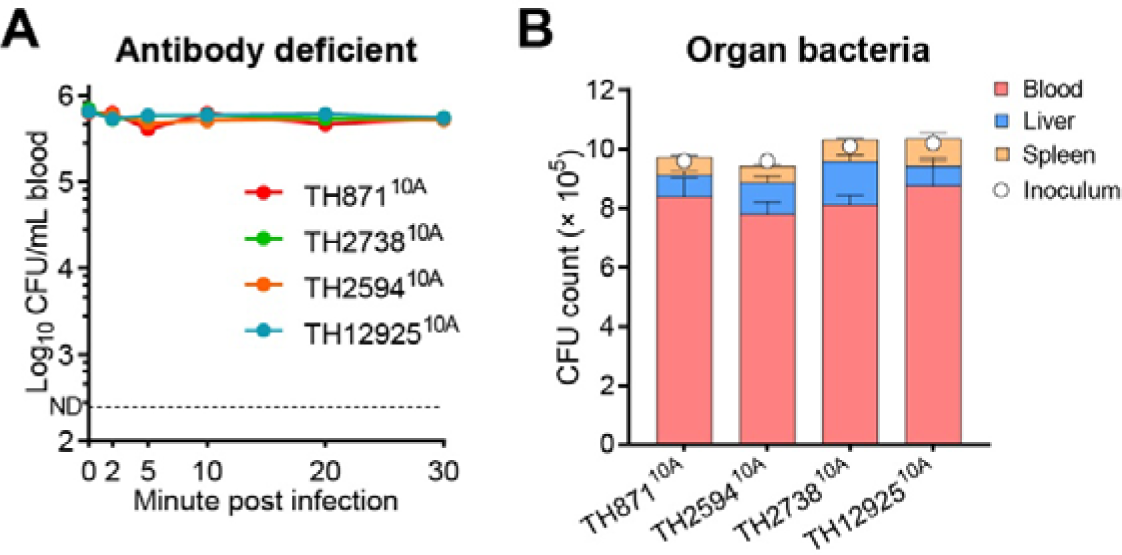
Clearance of other *Spn*10A strains in μMT mice. **(A** and **B)** Bacteremia kinetics in the first 30 min (A) and bacterial distribution of μMT mice at 30 min (B) post i.v. infection with 10^6^ CFU of four serotype-10A pneumococcal strains. n = 3.

**Figure S3.**
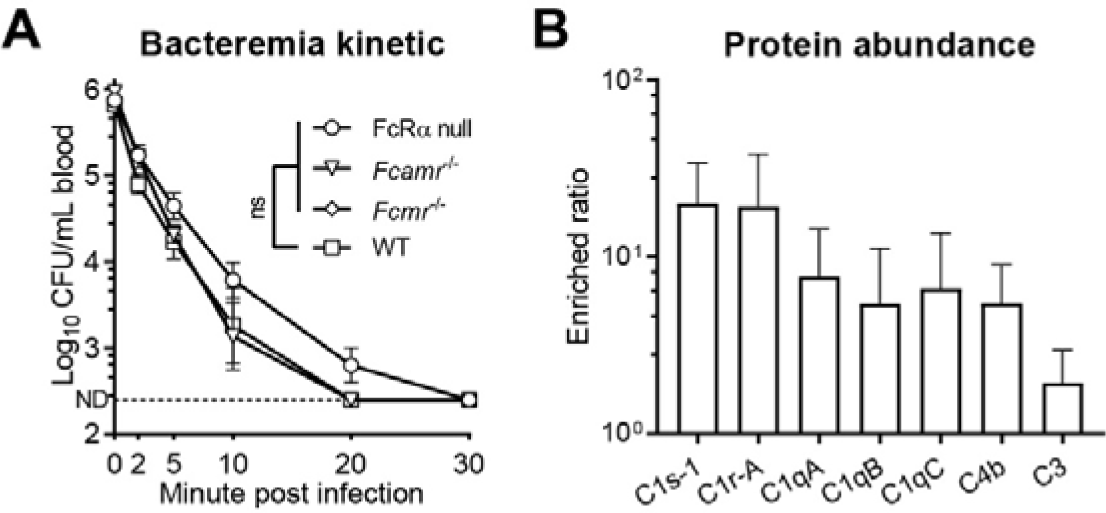
Dispensable role of Fc receptors in natural antibodies-mediated *Spn*10A capture. **(A)** Bacteremia kinetics of WT, FcRα null, *Fcmr*^-/-^ and *Fcamr*^-/-^ mice in the first 30 min post i.v. infection with 10^6^ CFU of TH860^10A^. n = 3. **(B)** Mass spectrometry results showing enriched complement components by CPS10A-beads compared to that of CPS10A Δ*wcrG*-beads. n = 3. The detailed results were listed in Table S1. Ordinary two-way ANOVA with Tukey’s multiple comparisons test (A) was performed. ns, not significant.

**Figure S4.**
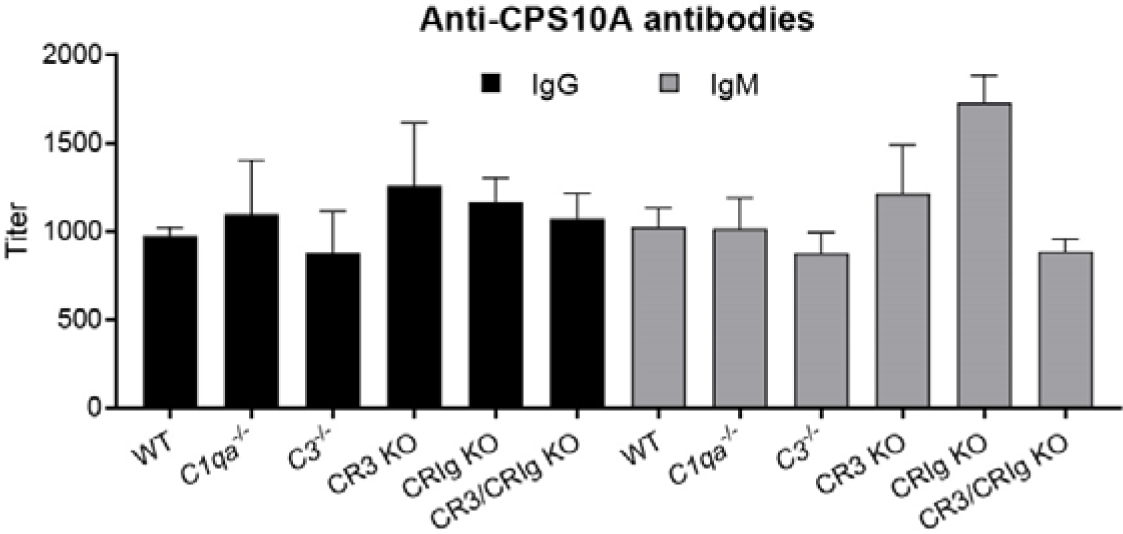
Comparable anti-CPS10A natural antibodies levels between WT and complement system-deficient mice. ELISA detection of IgG and IgM to CPS10A in serum of WT, *C1qa*^-/-^, *C3*^-/-^, CR3 KO, CRIg KO and CR3/CRIg KO mice. n = 5.

**Figure S5.**
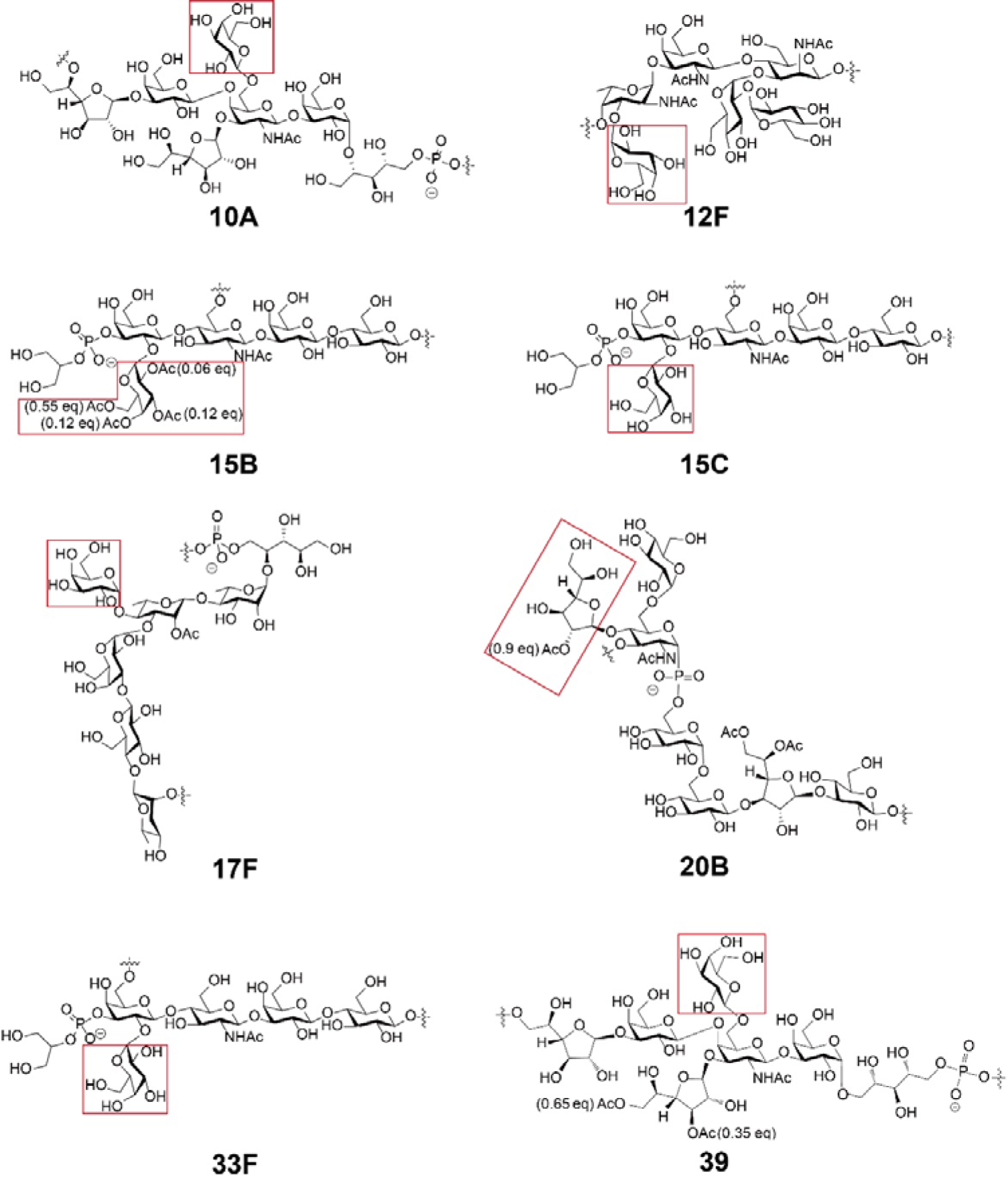
Structure of repeating unit of pneumococcal CPSs with galactose branch. Structure of repeating units of serotype-10A, -12F, -15B, -15C, -17F, -20B, -33F and -39 capsule were shown, in which galactose branches were labelled.

**Table S2.** Bacterial strains used in this study.

| **Strain ID** | **Species** | **Serotype** |
| --- | --- | --- |
| D39 | *S. pneumoniae* | 2 |
| TIGR4 | *S. pneumoniae* | 4 |
| TH860 | *S. pneumoniae* | 10A |
| TH860 $\Delta wcrG$ | *S. pneumoniae* | / |
| TH871 | *S. pneumoniae* | 10A |
| TH2594 | *S. pneumoniae* | 10A |
| TH2738 | *S. pneumoniae* | 10A |
| TH2912 | *S. pneumoniae* | 14 |
| TH12925 | *S. pneumoniae* | 10A |
| TH16770 | *S. pneumoniae* | 10B |
| TH16771 | *S. pneumoniae* | 10B |
| TH16772 | *S. pneumoniae* | 10B |
| TH16827 | *S. pneumoniae* | 39 |
| TH17033 | *K. pneumoniae* | K50 |

**Table S3.**
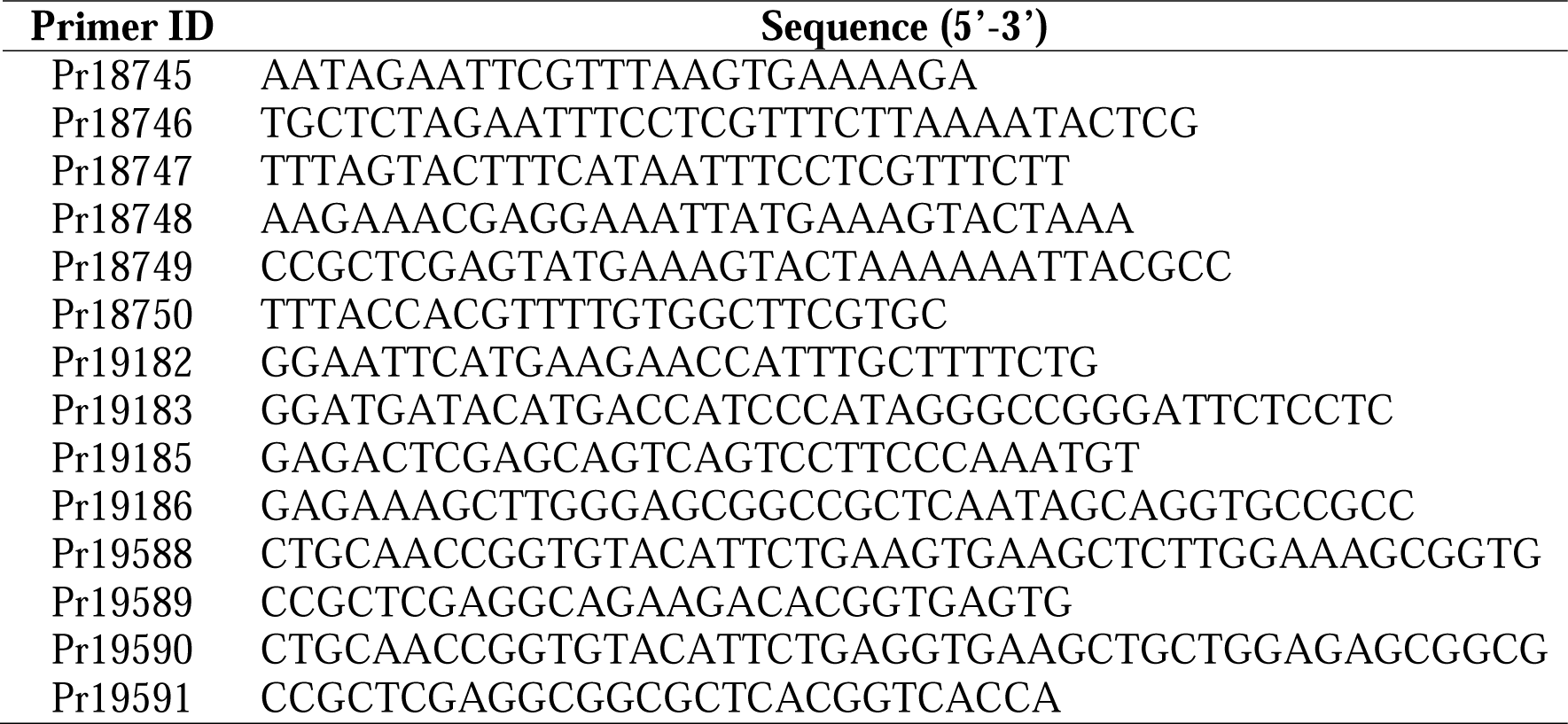
Primers used in this study.

**Table S4.** Construction of recombinant plasmids for antibody production.

| Plasmid ID | Backbone | Insertion segments | Primers for segments | Template | Digestion site |
| --- | --- | --- | --- | --- | --- |
| pTH17215 | pTH16770<br>(Wang et al., 2023) | VH X24 | / | Uniprot:<br>HVM39_MOUSE | AgeI/BsiWI |
| pTH17213 | pTH16771<br>(Wang et al., 2023) | VL X24 | / | Uniprot:<br>KV6A2_MOUSE | AgeI/XhoI |
| pTH17224 | Heavy chain<br>vector for<br>mouse IgM | J chain +<br>VH X24<br>+ CH<sub>IgM</sub> | Pr19182/Pr19183<br>(J) | Mouse spleen cDNA | EcoRI/HindIII |
|  |  |  | Pr19588/Pr19589<br>(VH X24) | pTH17215 |  |
|  |  |  | Pr19185/Pr19186<br>(CH<sub>IgM</sub>) | Mouse spleen cDNA |  |
| pTH17218 | pTH16770<br>(Wang et al., 2023) | VH X44 | / | Uniprot:<br>HVM37_MOUSE | AgeI/BsiWI |
| pTH17216 | pTH16771<br>(Wang et al., 2023) | VL X44 | / | Uniprot:<br>KV6A1_MOUSE | AgeI/XhoI |
| pTH17225 | Heavy chain<br>vector for<br>mouse IgM | J chain +<br>VH X44<br>+ CH<sub>IgM</sub> | Pr19182/Pr19183<br>(J) | Mouse spleen cDNA | EcoRI/HindIII |
|  |  |  | Pr19590/Pr19591<br>(VH X44) | pTH17218 |  |
|  |  |  | Pr19185/Pr19186<br>(CH<sub>IgM</sub>) | Mouse spleen cDNA |  |

**Video 1. Intravital microscopy (IVM) shows diminished liver capture of *Spn*10A in KC-deficient mice.** *S. pneumoniae* (green), KCs (red) and microvasculature (cyan) in the liver sinusoids of *Clec4f*-DTR mice treated with (+DT) or without (-DT) diphtheria toxin in the first 1 min post i.v. infection 5 × 10^7^ CFU of TH860^10A^ were shown. Graphic analysis was shown in Fig. 1E.

**Video 2. IVM shows inhibition of free CPS10A but not CPS10A without** β**1-6-linked galactose branch on KC capture of *Spn*10A.** Pneumococcal capture in the liver sinusoids of WT mice treated with 400 μg CPS10A or CPS10A Δ*wcrG* post i.v. infection 5 × 10^7^ CFU of TH860^10A^ were shown. Graphic analysis was shown in Fig. 2K.

**Video 3. IVM shows diminished KC capture of *Spn*10A in** μ**MT mice.** Pneumococcal capture in the liver sinusoids of WT or μMT mice post i.v. infection 5 × 10^7^ CFU of TH860^10A^ were shown. Graphic analysis was shown in Fig. 4F.

**Video 4. IVM shows unaffected KC capture of *Spn*14 in natural antibodies-deficient mice.** Pneumococcal capture in the liver sinusoids of WT, μMT or *Cd19*^-/-^ mice post i.v. infection 5 × 10^7^ CFU of TH2912^14^ were shown. Graphic analysis was shown in Figs. 4F and 4L.

**Video 5. IVM shows diminished KC capture of *Spn*10A in *Cd19*^-/-^ mice.** Pneumococcal capture in the liver sinusoids of WT or *Cd19*^-/-^ mice post i.v. infection 5 × 10^7^ CFU of TH860^10A^ were shown. Graphic analysis was shown in Fig. 4L.

**Video 6. IVM shows diminished KC capture of *Spn*10A in complement-deficient mice.** Pneumococcal capture in the liver sinusoids of WT or *Cd19*^-/-^ mice post i.v. infection 5 × 10^7^ CFU of TH860^10A^ were shown. Graphic analysis was shown in Fig. 5D.

**Video 7. IVM shows diminished KC capture of *Spn*10A in complement receptor-deficient mice.** Pneumococcal capture in the liver sinusoids of WT, CR3 KO, CRIg KO or CR3/CRIg KO mice post i.v. infection 5 × 10^7^ CFU of TH860^10A^ were shown. Graphic analysis was shown in Fig. 5J.

**Video 8. IVM shows diminished KC capture of *Spn*39 in natural antibodies-deficient mice.** Pneumococcal capture in the liver sinusoids of WT, μMT, *C3*^-/-^ or *C3*^-/-^ μMT mice post i.v. infection 5 × 10^7^ CFU of TH16827^39^ were shown. Graphic analysis was shown in Fig. 7D.

